# RNA synthesis and degradation regulate biomolecular condensates through non-equilibrium feedback

**DOI:** 10.64898/2026.05.11.724287

**Authors:** Ignacio Sanchez-Burgos, Andres R. Tejedor, Alberto Ocana, Jorge R. Espinosa, Rosana Collepardo-Guevara

## Abstract

Transcriptional condensates operate far from equilibrium, where continuous RNA synthesis and degradation dynamically reshape condensate composition. To investigate how RNA synthesis regulates condensate properties at sub-molecular resolution, we introduce **REACT-RNA**, a chemically specific coarsegrained molecular dynamics framework that explicitly couples RNA polymerisation, degradation, and nucleotide fluxes to sequence-dependent protein–RNA phase behaviour, and apply it to FUS and MED1 condensates. We find that RNA polymerization occurs preferentially at condensate interfaces because interfaces promote a highly reactive microenvironment that combines nucleotide influx, enrichment of reactive RNA chain ends, and enhanced molecular dynamics. Interfacial RNA polymerisation generates a spatially organised RNA population, with newly synthesised short RNAs enriched near the interface and longer RNAs partitioning towards the condensate core, which decreases the interfacial free energy and promotes multivalent RNA–protein interactions. As RNA concentration and chain length evolve over time, condensate composition, density, and stability change dynamically in response. Sustained RNA polymerisation drives re-entrant condensate dissolution, even of aged gel-like condensates, whereas RNA degradation stabilises long-lived non-equilibrium condensates containing elevated RNA content. Our results suggest that RNA synthesis, degradation, and nucleotide fluxes drive transcriptional condensates out of thermodynamic equilibrium while condensates in turn promote reactive molecular configurations that favour RNA production, enabling transient accumulation of RNA-rich compositional imbalance during bursts of transcription.

## I. INTRODUCTION

Biomolecular condensates in cells are embedded in a nonequilibrium intracellular environment. Some condensates compartmentalise biochemical reactions and energy-consuming processes, including transcription, RNA processing, and ATP-driven enzymatic activity^1,2^. These reactions continuously transform, create, and remove molecular components, thereby driving condensates away from thermo-dynamic equilibrium^3,4^. This suggests that biomolecular condensates not only regulate intracellular biochemistry^5–8^, but also continuously remodel their emergent properties— such as molecular organisation, stability, and viscoelastic behaviour—in response to their chemical activity. This coupling between biochemical activity and condensate biophysical properties supports the idea that cellular condensates behave as history-dependent non-equilibrium materials that are continuously shaped by their evolving composition.

Transcriptional condensates, which host transcription-associated chemical reactions, have been proposed to form near active genes, where they locally enrich transcription factors, coactivators such as Mediator (MED1), RNA-binding proteins including FUS, RNA Polymerase II, and ribonucleotide substrates^9–12^. By dynamically concentrating this machinery, such condensates provide environments that enhance local interaction probabilities and facilitate the assembly of the transcriptional apparatus^11,13^. Transcription has been suggested to occur near or at the interface of transcriptional condensates^11,13,14^. In this scenario, RNA Polymerase II polymerises free ribonucleotides into nascent RNA chains that elongate over time, while degradation, processing, and export limit RNA accumulation^13^.

Increasing evidence indicates that nascent RNA is not only a product of transcriptional condensates but also an active regulator of their formation, stability, and other biophysical properties^9–11,15–17^. Most current understanding of this behaviour, however, is based on systems in which RNA concentration and length are static variables. In solutions of RNA-interacting proteins, including HP1α^18^, G3BP1^5^, and FUS^17,19^, RNA at moderate concentrations—approaching charge neutrality—promotes phase separation by acting as a multivalent scaffold that increases condensed-phase network connectivity^20–23^. However, at high concentrations— typically beyond electroneutrality—RNA instead drives condensate dissolution^5,11,16,22,24^, revealing a re-entrant dependence of condensate stability on RNA concentration^25^. RNA concentration is therefore a key control parameter governing RNA–protein condensates^26^. Beyond concentration, RNA length further modulates condensate stability, particularly when protein–RNA interactions dominate network connectivity^22^.

In transcriptional condensates, however, both RNA length and concentration change dynamically, providing a time-dependent source of material flux that drives these condensates away from thermodynamic equilibrium. As a result, their composition—and arguably their internal molecular organisation and material properties—continuously evolve over time rather than remaining static. RNA accumulation and elongation initially promote condensate assembly but ultimately drive destabilisation at high concentrations. This re-entrant behaviour has been proposed to underlie transcriptional bursting, whereby low levels of nascent RNA stabilise condensates and enhance transcriptional activity, while subsequent RNA accumulation feeds back to destabilise the condensate and terminate the burst, thereby establishing a non-equilibrium feedback loop driven by continuous RNA production^13^.

Despite this emerging picture, how time-dependent RNA growth and degradation control condensate stability, material properties, and internal organisation at sub-molecular resolution remains unclear. This limitation arises in part because experiments cannot yet directly resolve the coupled time-dependent evolution of molecular interactions, RNA length, and concentration within condensates to the regulation of their mesoscale properties. Molecular simulations provide a complementary approach by enabling direct control over these processes at molecular resolution. Although inherently approximate, molecular simulations allow individual mechanisms—such as RNA growth, degradation, and interaction strengths—to be systematically tuned and isolated. Most existing simulation frameworks, however, treat RNA length and concentration as fixed parameters, which does not capture the dynamic coupling between reactions and condensate behaviour.

In this work, we develop **REACT-RNA** (Reactive RNA polymerisation and degradation), a non-equilibrium residue and nucleotide-resolution coarse-grained molecular dynamics (MD) framework that explicitly couples RNA polymerisation and degradation to the condensate and its evolving properties. RNA polymerisation is implemented as a dynamic, local bond-formation process, with reaction events conditioned by the surrounding protein environment. REACT-RNA is built upon the Mpipi-Recharged model^27^, enabling investigation of reaction-driven changes in RNA length and concentration with sequence specificity. This framework moves beyond equilibrium simulations of protein–RNA condensates, in which RNA length and concentration are fixed parameters, and instead enables simulations of condensates with time-dependent RNA compositions set by the balance of RNA production and degradation rates, and free nucleotide fluxes.

REACT-RNA, summarised in Figure 1(a), includes three key features: (i) dynamic formation of inter-nucleotide (phosphodiester) bonds constrained to protein-crowded environments, (ii) coupling of RNA chain growth to an explicit nucleotide reservoir, thereby enabling sustained fluxes of matter^28,29^, and (iii) RNA degradation as a competing process. This unified framework captures the nonequilibrium degradation of RNA and its coupling to protein–RNA interaction networks.

**Figure 1:**
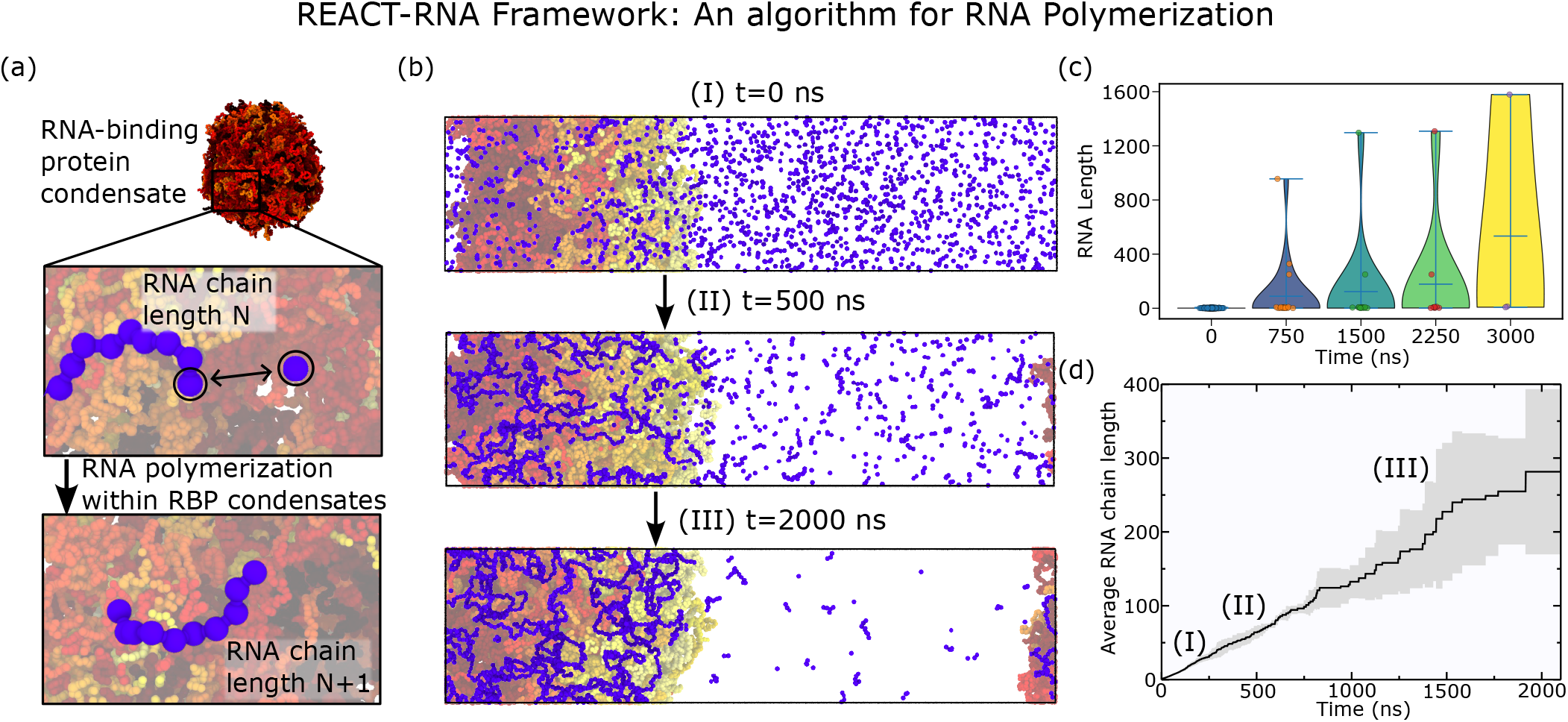
Representative simulation in which RNA molecules emerge and grow within FUS condensates. (a) Illustrative rendering of an FUS protein condensate, showing RNA molecules (blue) undergoing growth within a protein-crowded environment (red). In this simulation we allow RNA polymerization in protein-crowded environments, but no turnover or exchange with a precursor reservoir is enabled. This simulation was performed at 290 K, which is below the critical temperature for FUS phase separation (see Figure 2(a)). (b) Time evolution of the RNA polymerization process in presence of a pre-formed FUS condensate at T = 287.5 K (below the critical temperature for phase separation). (c) Time-resolved RNA length distributions from violin plots (d) Average RNA strand length as a function of time during the RNA growth example simulation. Data is averaged from 3 independent trajectories, and the gray shade represents the uncertainty, estimated as the standard deviation.

Using REACT-RNA, we investigate how RNA synthesis and degradation reshape the phase behaviour of FUS and MED1 condensates under nonequilibrium conditions. We show that ongoing RNA growth dynamically reorgan-ises condensates by altering RNA length distributions, intermolecular connectivity, and electrostatic interactions, giving rise to concentration-dependent stabilisation and re-entrant dissolution. RNA production becomes spatially coupled to condensate interfaces, linking reaction dynamics to molecular exchange and generating history-dependent condensate organisation. Simultaneously, condensates favour reactive molecular encounters that promote RNA production. We further show that RNA degradation enables condensates to transiently sustain excess RNA and negative charge beyond equilibrium electroneutrality constraints. Together, our results establish a molecular framework for understanding how transcription-associated RNA growth, degradation and fluxes dynamically regulate the formation, stability, and material properties of biomolecular condensates.

## II. RESULTS AND DISCUSSION

### A. REACT-RNA: A Molecular Dynamics Framework for Time-Dependent RNA Polymerization in Sequence-specific Condensates

Understanding how RNA polymerisation and degradation regulate condensate formation and properties requires an explicit treatment of time-dependent changes in both RNA length and concentration. Equilibrium descriptions capture only static snapshots of this process—for example, how fixed RNA lengths and concentrations affect condensate properties—and therefore provide an incomplete picture. In addition, because polymerisation and degradation dynamically create, remove, and rearrange intermolecular contacts, they break detailed balance and introduce history dependence into the dynamics of condensates.

To investigate this problem with chemical specificity, we develop **REACT-RNA**, a non-equilibrium MD framework that explicitly treats RNA polymerisation and degradation as time-dependent processes occurring within sequence-specific protein–RNA condensates. This framework is implemented within the Mpipi-Recharged coarse-grained model for biomolecular phase separation^27^, which represents amino acids and nucleotides at residue resolution (numerical details, force-field potentials, and parameters are provided in the Supplementary Material (SM), Section SI). This integration enables RNA polymerisation dynamics to be resolved directly at the level of sequence-specific intermolecular interactions.

Mpipi-Recharged considers pair-specific parameters for both ionic and non-ionic interactions, and therefore captures experimentally observed behaviour of highly charged biomolecular condensates, including charge patterning and stoichiometric effects in protein–RNA condensates without requiring explicit solvent^27^. It also reproduces the behaviour of a broad range of protein sequences, including condensate phase diagrams, variations in viscoelastic properties as a function of sequence variations, as well as single-protein radii of gyration as a function of temperature and salt concentration^30–35^. In addition, Mpipi-Recharged enables sim-ulations of system sizes sufficient to observe the collective assembly of tens to hundreds of proteins during timescales spanning several microseconds^23,33^.

Motivated by evidence that RNA synthesis is spatially coupled to the assembly of transcriptional condensates, which enrich the transcriptional machinery together with its molecular substrates^9,10,12–14^, we implement in REACT-RNA a sequential nucleotide–nucleotide bond formation mechanism between terminal RNA beads and free nucleotides in solution using an on-the-fly bond-formation algorithm (see SM)^36,37^. In this framework, reaction events are further modulated by the local condensate environment, allowing RNA chain elongation to respond dynamically to spatial variations in condensate composition, structure, and molecular crowding. Specifically, bonds form only when (i) two terminal nucleotides are found within a cut-off distance of 7.5 Å, and (ii) each nucleotide is simultaneously in contact with at least one protein residue within a 10 Å cut-off. These criteria restrict polymerisation to protein-rich environments and couple RNA growth to the local interaction network (see SM section SXIII for a detailed discussion on this cut-off values choice).

Our approach captures the minimal physicochemical definition of transcription as the polymerisation of ribonucleotides into RNA^38^, without explicitly modelling the en-zymatic action of RNA Polymerase II. This allows us to investigate how condensate properties regulate RNA production, and conversely, how RNA growth reshapes condensate structure, connectivity, and stability. Within this framework, RNA length and concentration emerge as dynamical variables rather than externally imposed parameters.

Further implementation details and input scripts are provided in the SM (Section SIV). Figure 1(b) illustrates this process for a FUS–RNA condensate at different times (no RNA turnover or precursor reservoir is introduced here), showing that RNA growth occurs within the condensed phase. Figures 1(c) and (d) show the time evolution of RNA strand length and the average chain length, respectively.

### B. RNA polymerisation Stabilises FUS Condensates under Limited Nucleotide Supply

In this section, we use REACT-RNA to investigate how FUS condensates respond to a minimal non-equilibrium scenario in which RNA polymerisation occurs in a closed system containing a finite solution of nucleotide precursors and in the absence of RNA degradation. This setup isolates the effect of time-dependent RNA growth on condensate stabil-ity, while avoiding additional contributions from molecular degradation or exchange with an external reservoir. As a result, RNA length changes dynamically but the total amount of nucleotides remains bounded. The system therefore approaches a well-defined state as the finite nucleotide pool is progressively consumed. In following sections, we will extend the framework to include nucleotide exchange and RNA degradation.

We now investigate how RNA polymerisation affects the formation of FUS condensates under conditions where the system does not undergo phase separation at equilibrium in our model. For this, we first determine the equilibrium phase diagram of FUS using standard direct coexistence simulations in the temperature–density plane with the Mpipi-Recharged model (Figure 2(a)). We then perform REACT-RNA simulations at the critical solution temperature identified (Figure 2(a), i.e. 337 K) in systems containing fixed amounts of FUS and ribonucleotides. Specifically, we initialise a well-mixed system containing FUS and a limited pool of uridine nucleotide precursors and then activate the RNA polymerisation framework.

**Figure 2:**
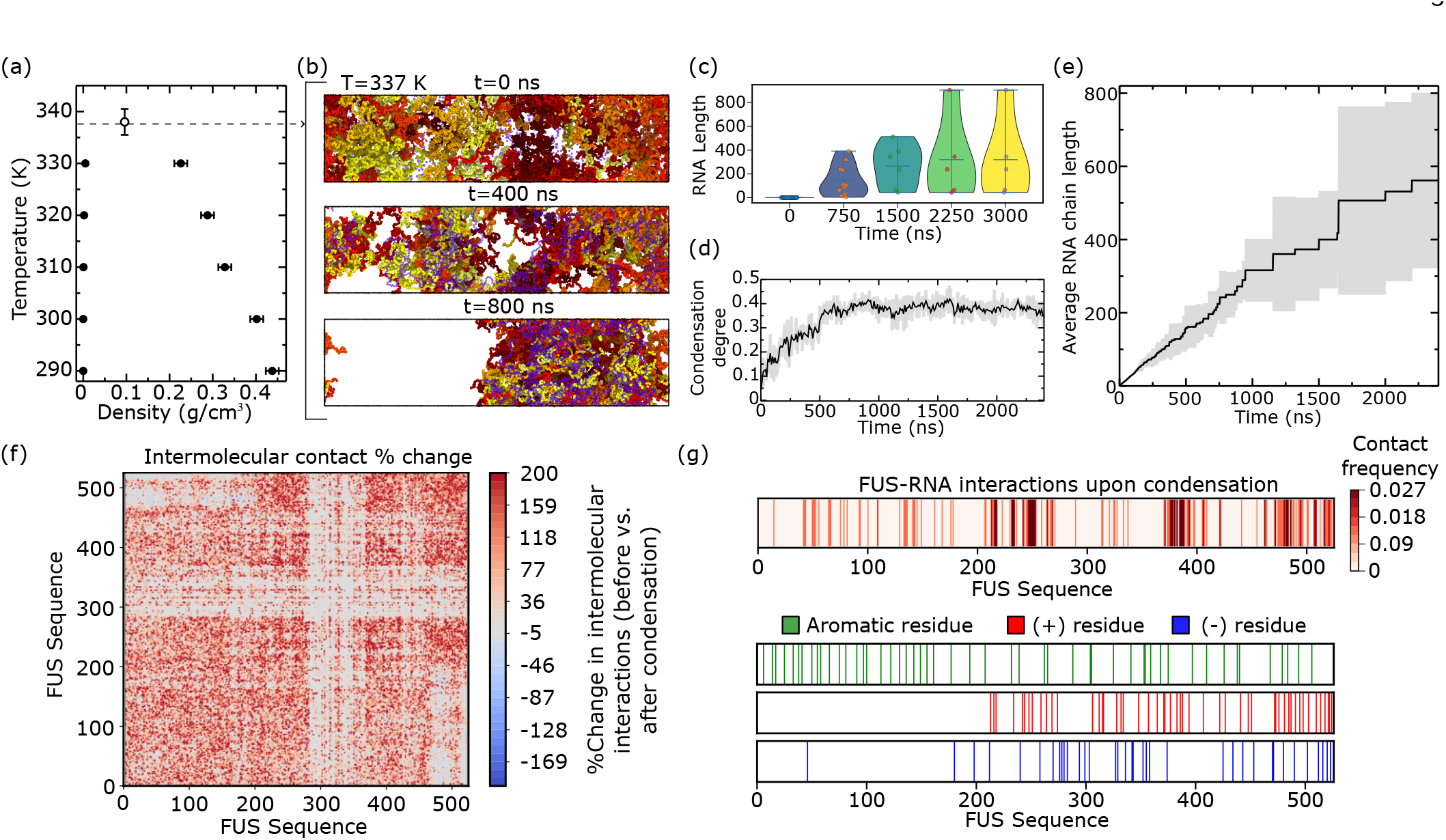
Temperature-density phase diagram of FUS. The filled dots represent densities directly measured from direct coexistence simulations, while the empty point depicts the critical point, determined through the law of rectilinear diameters and critical exponents (see SM section SIII for more details on this calculation). (b) Rendered images depicting the condensation process at different times. The FUS molecules are colored in red tones while the RNA molecules are depicted with faded blue for better condensate visualization. This simulation was performed under critical conditions T = 337 K. (c) Time-resolved RNA length distributions from violin plots. (d) Normalized condensation degree parameter over time. The shaded area represents the uncertainty, estimated as the standard deviation between the 3 replicas simulated. (e) Average RNA chain length over time during the RNA polymerization simulation. The shaded area represents the uncertainty, estimated as the standard deviation between the 3 replicas simulated. (f) Intermolecular contact map difference for FUS-FUS intermolecular interactions expressed in %. The initial and final steps are t=0 ns (no condensation) and t=1000 ns (condensate formed). Additional simulations with no RNA polymerization were performed for this calculation. (g) Top: FUS-RNA intermolecular contacts frequency map measured upon condensation (t=1000 ns). Bottom: FUS sequence residue map where the aromatic (green), positive (red) and negatively charged (blue) residues are highlighted.

As shown in Figure 2(b), condensate formation emerges only after RNA elongation, demonstrating that RNA polymerisation stabilises FUS condensates under near-critical conditions where the protein alone does not phase separate at equilibrium. Analysis of the time evolution of RNA length distributions (Figure 2(c)) shows that condensation is triggered once RNA reaches lengths sufficient to engage multiple FUS molecules simultaneously, i.e. exceeding ∼ 150 nucleotides for poly-uridine (poly-U) strands. Notably, RNA length distributions remain broad over time (Figure 2(c)), indicating that condensation is sustained by a heterogeneous population of RNA chains of different lengths. The subpopulation of longer RNAs increases the inter-molecular network connectivity of the condensate, while shorter chains contribute more weakly to the network, but can stabilise the interfaces^23^. Importantly, because RNA growth is limited by a finite pool of nucleotide precursors, RNA length saturates over time. The system therefore approaches a steady state in which RNA-mediated multivalency is sufficient to stabilise a condensed phase without entering the high-RNA-concentration regime associated with re-entrant dissolution.

This behaviour arises from a feedback mechanism where condensate properties and reaction kinetics are coupled. That is, increased local protein density enhances RNA polymerisation and global concentration (within the proteincoupled reaction mechanism implemented here) by promoting nucleotide encounters and stabilising reactive configurations, while RNA elongation increases multivalency and strengthens the protein intermolecular network connectivity.

To quantify condensate formation, we define a condensation degree parameter based on the spatial distribution of protein density. The simulation box is partitioned into slabs along its elongated axis, and each slab is classified as belonging to the condensed phase if its local density exceeds 25% of the maximum density across all slabs (see section SIX for details on parameter threshold optimization). The condensation degree is then defined as the fraction of such condensed slabs relative to the total number of slabs and normalized between 0 and 1 to enable observing system evolution and comparison across simulations (Figure 2(d)). Consistently, the time evolution of the condensation degree (Figure 2(d)) shows that phase separation is initiated at early times, when RNA chains are still short, whereas the subsequent superlinear increase in RNA length (Figure 2(e)) indicates that polymerisation kinetics accelerate as the condensate becomes denser. As a result, the RNA polymerisation reaction, sustained by nucleotide consumption, maintains the system in a non-equilibrium regime in which a condensate is stabilised under conditions where phase separation is not observed in the absence of RNA polymerisation. In SM section SXII, we demonstrate that RNA polymerization is essential for the condensation process by comparing it with a system in which the REACT-RNA algorithm is disabled. In the absence of RNA polymerization, the system remains in a single dilute phase, and no condensation is observed. Because the system operates with a finite pool of nucleotide precursors, polymerisation eventually ceases, and the system relaxes towards an equilibrium state at long times. Importantly, the resulting equilibrium state is not unique but history dependent. Indeed, the distribution of RNA chain lengths, once the nucleotide pool is exhausted, is effectively frozen in the absence of degradation, leading to a condensate composition that reflects the prior nonequilibrium dynamics. This behavior is consistent with previous *in vitro* studies^16,17,39^, showing that RNA can act as a multivalent scaffold, enhancing intermolecular connectivity and effectively lowering the threshold for condensate nucleation^21^.

To elucidate the molecular basis of RNA-induced condensation, we analyse the change in FUS–FUS intermolecular contacts between the initial state (t = 0 ns, homogeneous phase) and the final state (t = 1000 ns, fully developed condensate following RNA growth) (Figure 2(f)). We define an effective contact when two residues are closer than 1.2 σ, where σ is the average molecular diameter of the residue pair (see SM section SX for a detailed anaylsis of this cutoff distance optimization). We observe a global increase in intermolecular contacts across the FUS sequence, consistent with the formation of a dense, percolated interconnected protein network^20^. This increase is spatially heterogeneous along the sequence (Figure 2(f)), indicating that RNA-mediated condensation selectively amplifies interactions in specific regions rather than uniformly across the protein. Notably, this increase is less pronounced within the RNA recognition motif (residues 288–371), a structured globular domain with reduced capacity to form the weak, transient interactions that stabilize liquid-like condensates. This observation is in line with previous studies indicating that intrinsically disordered regions, rather than folded domains, primarily drive condensate formation^40–44^.

We further characterize protein–RNA coupling by examining FUS–RNA interaction profiles (Figure 2(g), top), alongside a residue-level map highlighting aromatic, positively charged, and negatively charged amino acids (Figure 2(g), bottom). This analysis reveals a sequence-dependent binding pattern, with contacts strongly enriched in regions containing positively charged residues, particularly arginine and lysine. These interactions are consistent with electrostatic attraction between the negatively charged RNA backbone and cationic protein domains, in agreement with previous studies of RNA–binding protein condensates^24,45^.

### C. RNA polymerisation Drives Re-entrant MED1 Coacervation Under Sustained Nucleotide Supply

Having established that RNA polymerisation stabilises FUS condensates under near-critical conditions, we next asked whether similar feedback mechanisms emerge in systems whose phase separation is driven by heterotypic protein–RNA interactions rather than homotypic protein interactions. To address this question, we examined MED1, a transcriptional coactivator whose IDRs exhibit limited ability to phase separate via homotypic interactions but readily undergo phase separation in the presence of nucleic acids.

MED1 plays a central role in transcriptional condensates by linking DNA-associated factors with RNA polymerase II^46^. Following Henninger *et al*.^11^, we model the MED1 IDR, which is characterized by a low content of aromatic residues and a high density of positively charged residues (see sequence in SM section SII). In agreement with previous equilibrium simulations with static compositions using the CALVADOS force field^24^, the Mpipi-Recharged model predicts that this region does not undergo intrinsic phase separation at physiological or near-physiological temperatures.

We then extend REACT-RNA to include continuous ex-change with a surrounding reservoir of nucleotide precursors. In this regime, RNA length and concentration are no longer intrinsically bounded, as the nucleotide reservoir allows persistent RNA polymerisation and continuous molecular flux into the condensate. Specifically, we maintain a constant nucleotide concentration in the dilute phase by introducing a local order parameter that monitors nucleotide density and periodically performs insertion or deletion moves (every 5 ns). This nucleotide concentration includes all nucleotides, either those that are part of a longer RNA chain or single precursors. Therefore, with this framework, under RNA excess both free nucleotides as well as polymerized molecules completely embedded in the diluted phase are subject to deletion, while only single nucleotides are inserted in RNA shortage conditions. Further details on this algorithm are provided in SM section SV, and a detailed discussion on the reservoir concentration in SM section SXI.

We use REACT-RNA with the nucleotide reservoir to examine how a solution of MED1 IDRs responds to sustained RNA polymerization. Starting from a well-mixed mixture of MED1 and nucleotides (Figure 3(a), top), RNA polymerisation progressively generates RNA chains that elongate and introduce multivalent binding sites for MED1. This drives the emergence of a multi-component MED1–RNA condensate after ∼400 ns (Figure 3(a), middle panel). However, continued RNA polymerization and elongation, enabled by the flux of nucleotides entering the condensate from the diluted phase, alter the balance of interactions within the dense phase. As RNA accumulates and lengthens, MED1–RNA interactions increasingly saturate, and the effective RNA–RNA repulsive interactions encoded in the Mpipi-Recharged model become more prominent, weakening the effective intermolecular connectivity within the condensate. As a result, the condensate becomes progressively destabilised and ultimately dissolves at longer times (Figure 3(a), bottom). This behaviour is quantified by the normalised condensation degree (Figure 3(b)), which exhibits a re-entrant dependence on RNA production. Condensation is maximised at intermediate RNA levels, whereas continued RNA accumulation suppresses phase separation at long times (*t >* 600 ns). Additionally, in SM section SXIV we show the time evolution of the RNA content, the average chain length and the density profiles of the different components, further proving the association and relation between RNA growth, concentration and the re-entrant condensation process.

**Figure 3:**
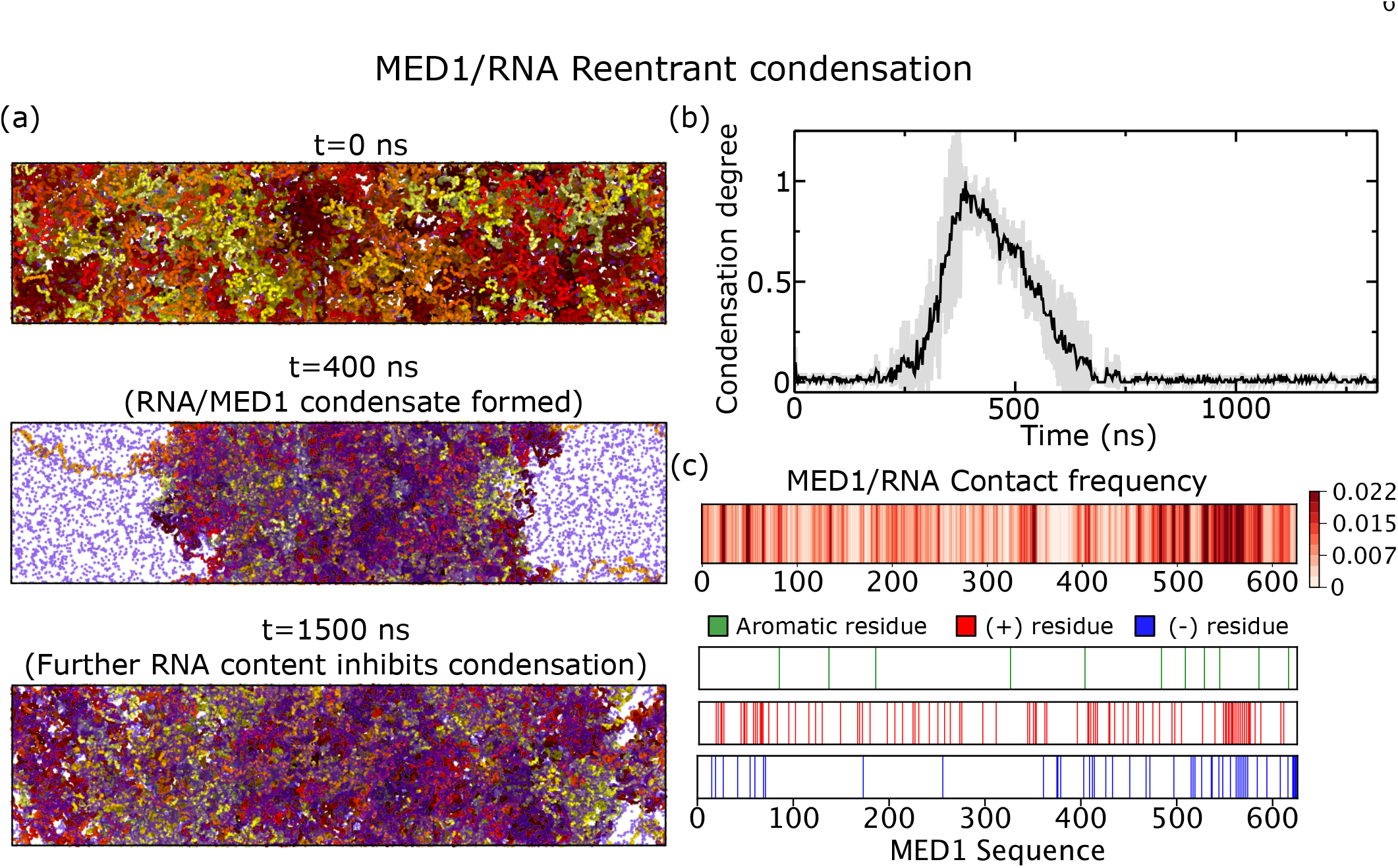
(a) Rendered images of the representative stages in the MED1/RNA transcriptional feedback loop. Top: No condensation before RNA growth. Middle: Condensation driven by RNA growth. Bottom: Excess of RNA hinders phase separation. (b) Normalized condensation degree parameter as a function of time. The shaded area represents the uncertainty of this quantity, estimated as the standard deviation between 3 independent simulation replicas. (c) Top: MED1–RNA intermolecular contacts frequency along the MED1 sequence. Bottom: MED1 sequence residue map where the aromatic (green), positive (red) and negatively charged (blue) residues are highlighted.

These results demonstrate that regulation of condensate stability by RNA polymerisation extends beyond homotypic protein condensates and also applies to systems assembled through heterotypic protein–RNA interactions and complex coacervation. More generally, they show that sustained RNA production in the absence of degradation can drive condensates into a re-entrant dissolution regime through progressive RNA accumulation, which in the Mpipi-Recharged model alters the balance of intermolecular interactions and ultimately destabilises the condensed phase.

Finally, we examine the molecular determinants of MED1–RNA interactions during this transient condensation regime. To this end, we computed the MED1–RNA inter-molecular contact map at *t* = 400 ns (Figure 3(c), top), corresponding to an intermediate state in which polymerisation was halted to isolate the underlying interaction patterns. In this map, colour intensity reflects the frequency of contacts between RNA and individual MED1 residues. This map is shown alongside the sequence distribution of aromatic, positively charged, and negatively charged residues (Figure 3(c), bottom), enabling a direct comparison between sequence composition and interaction propensity.

MED1–RNA interactions are dominated by positively charged residues, primarily arginine and lysine, which form favourable electrostatic interactions with the negatively charged RNA. These interactions are broadly distributed along the MED1 sequence, with enhanced binding in regions enriched in positive charge (e.g. residues 550–575). In contrast, negatively charged MED1 regions show reduced interaction frequencies with RNA. Therefore, the MED1– RNA interactions that drive condensation are largely electrostatic associations.

### D. Balance between RNA polymerisation and Degradation Controls the Stability and Composition of Non-equilibrium Condensates

Cellular systems operate under conditions where RNA concentration is dynamically regulated by a balance between synthesis and degradation. RNA synthesis is achieved, for instance, by RNA polymerase II within transcriptional condensates, whereas degradation processes mediated by RNases and RNA exosome complexes are essential for maintaining RNA homeostasis^10,47,48^. To capture this competition, in this section, we extend our REACT-RNA framework by incorporating RNA degradation as a stochastic removal process, in addition to the nucleotide reservoir. Specifically, at regular intervals of 5 ns, RNA chains are removed stochastically. Opposed to the RNA reservoir described in the previous section, his coarse-grained implementation mimics RNA degradation and limits unbounded polymer growth, targeting all molecules present in the simulation and not only those in the diluted phase. Details are provided in SM Section SVI.

To disentangle the role of RNA degradation, we first examine the consequences of sustained RNA growth in FUS condensates in the absence of degradation, before intro-ducing RNA degradation as a competing process. Starting from an initially well-mixed system at the critical temperature of FUS solutions, RNA polymerisation in the presence of a nucleotide reservoir initially induces condensate formation (Figure 4(a), top and middle), consistent with the multivalent scaffold role of RNA discussed in the previous sections. However, rather than stabilising a persistent condensate, excessive RNA accumulation disrupts the dense phase, producing smaller and less cohesive clusters (Figure 4(a), bottom). Therefore, as in the case of MED1, unbounded RNA synthesis initially promotes condensate assembly, but continued RNA production without degradation eventually drives the system into the re-entrant dissolution regime. This behaviour reflects the saturation of FUS–RNA interactions and the increasing contribution of RNA–RNA electrostatic repulsion at high RNA concentrations, which weakens the inter-molecular connectivity of the network required for condensate integrity. Such destabilisation is consistent with previous observations of a feedback loop in which RNA production initially promotes condensation but ultimately drives dissolution upon RNA overaccumulation^11,16^.

**Figure 4:**
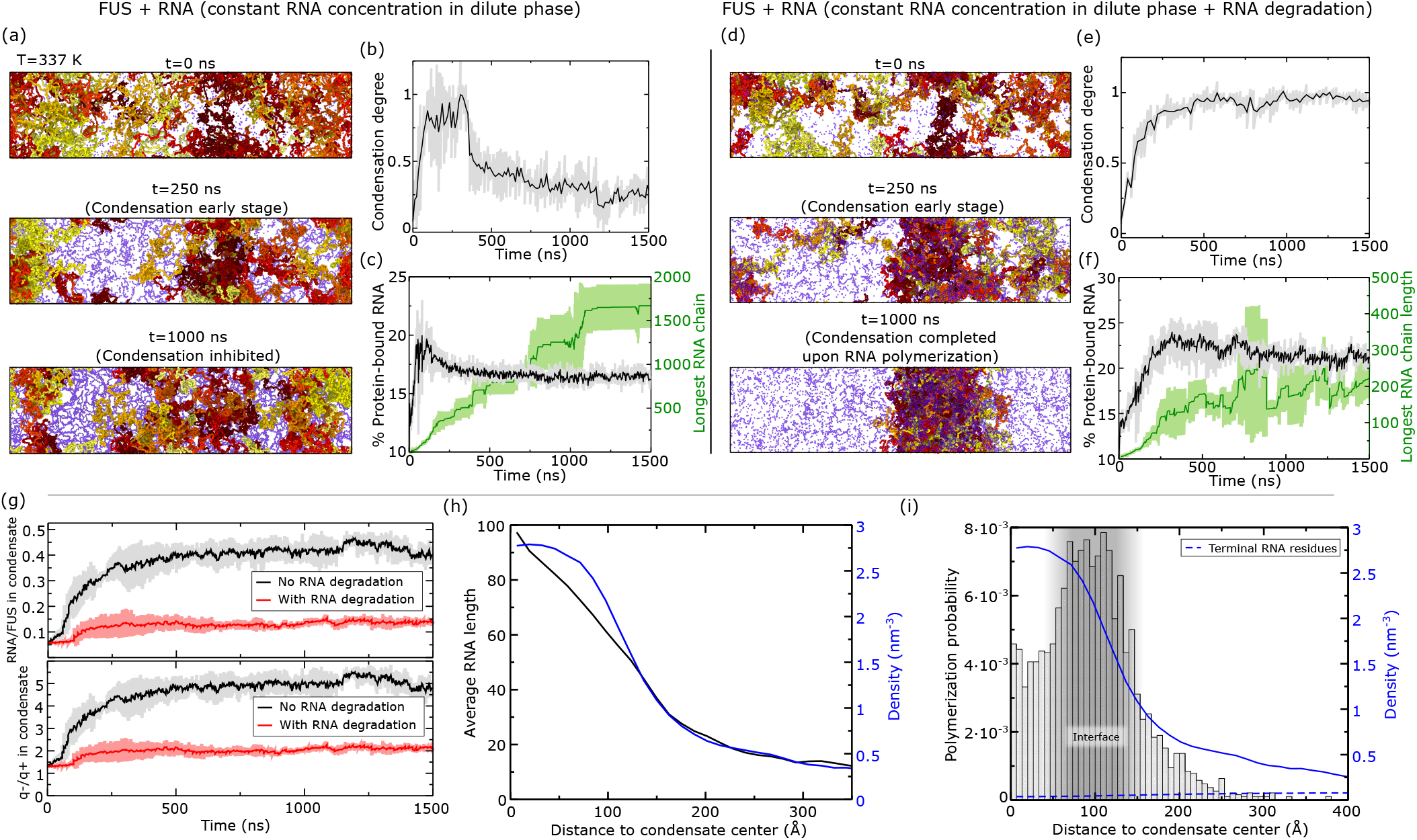
(a) Rendered images of the representative stages in the FUS/RNA feedback loop process. This simulation exchanges particles with a nucleotide reservoir, keeping the RNA concentration in the dilute phase constant. Top: No condensate prior to RNA growth. Middle: Condensation triggered by FUS-RNA interactions. Bottom: Condensation is not completed due to RNA overexpression. (b) Time dependence of the normalized condensation degree parameter for the simulation depicted in (a). For all panels in this Figure, the shaded area represents the uncertainty, estimated as the standard deviation between three independent simulation trajectories. (c) % of protein bound nucleotides in the system as a function of time, represented along the longest RNA chain present in the simulation depicted in (a). (d) Rendered images of the RNA polymerization-drive condensation. As in (a), this simulation exchanges particles with a nucleotide reservoir, but in this one we also implement the RNA degradation algorithm. Top: No condensation prior to RNA growth. Middle: RNA polymerization drives condensation. The FUS/RNA condensate remains stable when RNA molecules are growth-limited by the degradation process. (e) Time dependence of the normalized condensation degree parameter for the simulation depicted in (d). (f) % of protein bound nucleotides in the system as a function of time, represented along the longest RNA chain present in the simulation depicted in (d). (g) Top: RNA/FUS residue ratio (RNA nucleotides per FUS amino acid) time evolution measured in regions of the simulation box classified as the condensed phase, using the same relative density criterion applied for the condensation degree parameter. Results are shown for simulations with and without RNA degradation, as indicated in the legend. Bottom: Negative/positive charge ratio time evolution measured in regions of the simulation box classified as the condensed phase. Results are shown for simulations with and without RNA degradation, as indicated in the legend. (h) Average RNA chain length against the distance to the condensate center (black curve), represented alongside the density profile (blue curve). (i) Normalized polymerization probability distribution as a function of the distance to the condensate center, plotted alongside the density profile (blue curve) and terminal RNA beads (including single nucleotides, dashed blue curve).

We quantify this transition using the condensation degree parameter (Figure 4(b)), which increases rapidly at early times, reaching a maximum at ∼250 ns, and then decreases gradually, indicating destabilisation of the dense phase. To identify the molecular origin of condensate dissolution, we analyse the fraction of protein-bound RNA nucleotides together with the length of the longest RNA chain (Figure 4(c)). The fraction of protein-bound nucleotides initially rises sharply, consistent with enhanced inter-molecular interactions driving condensate assembly, but subsequently decreases as RNA over-accumulation weakens such interactions. Once FUS binding sites become saturated, additional RNA disrupts the network via increased RNA–RNA repulsion and suppressed RNA-mediated FUS–FUS bridging. Despite this, the longest RNA chain lengthens monotonically, reflecting continuous and irreversible RNA growth inside the remaining smaller FUS clusters. Thus, RNA polymerisation first increases the inter-molecular connectivity and promotes condensation, but eventually drives the system into re-entrant dissolution.

We next activate the RNA degradation algorithm to determine whether this can counterbalance sustained RNA production and stabilise the condensate. For a representative degradation probability of 0.1%, RNA-driven condensation still emerges as polymerisation proceeds (Figure 4(d)). Crucially, in contrast to the degradation-free case, dynamic degradation balances RNA growth and prevents excessive RNA accumulation. The system therefore evolves towards a non-equilibrium steady state at late times in which a condensate persists. This behaviour is quantified by the normalised condensation degree parameter (Figure 4(e)), which keeps increasing up to *t* ∼ 500 ns and subsequently plateaus, indicating the emergence of the persistent con-densed phase. We note that the degradation probability of 0.1% is presented here as an illustrative parameter choice. For sufficiently low degradation probabilities, RNA removal cannot compensate for sustained polymerisation, resulting in RNA accumulation within the condensate and its eventual dissolution. The full dependence of the condensate stability and dynamics on the degradation probability is discussed in Section SVI.

The time evolution of protein-bound RNA and the longest RNA chain further shows how degradation selects a stable dynamical regime (Figure 4(f)). In contrast to the degradation-free case (Figure 4(c)), the fraction of protein-bound RNA remains elevated and does not exhibit a pronounced decline, consistent with persistent condensate integrity. Meanwhile, the longest RNA chain fluctuates between ∼200 and 300 nucleotides, with intermittent drops arising from degradation events. This dynamically maintained length range is notable because previous equilibrium simulations showed that FUS phase separation is enhanced most efficiently by RNAs approaching or exceeding the characteristic size of the protein, which can bridge multiple FUS molecules and maximise multivalent connectivity^23^. Such RNA lengths have also been shown to efficiently bridge independent FUS proteins and drive phase separation upon binding^22,49^. Here, this optimal range is not imposed externally, but emerges dynamically from the RNA polymerisation– degradation balance. That is, when sustained RNA polymerisation is available, RNA degradation dynamically selects RNA lengths that maximise intermolecular connectivity while suppressing the overgrowth regime associated with re-entrant condensate dissolution.

We next examine how this dynamical regulation affects condensate composition and intra-molecular organisation. The RNA/FUS ratio within the condensed phase (Figure 4(g)(top)) is markedly higher in the degradation-free simulation than in the degradation-controlled condensate, consistent with progressive RNA accumulation preceding condensate dissolution in our model. To further characterise condensate composition, we analysed the ratio of negatively charged RNA nucleotides to positively charged protein residues within the condensed phase (Figure 4(g); bottom). While this quantity provides a useful measure of the relative abundance of RNA and positively charged protein residues, it should not be interpreted as a direct measure of the net charge of the condensate. The total charge of a biomolecular condensate is determined jointly by its biomolecular components and the partitioning of mobile ions^50^, and must satisfy overall electroneutrality. Thus, even condensates in which the biomolecular components alone carry a non-zero net charge might remain close to electroneutrality through the partitioning of counterions^51^, which are only implicitly represented in our model. Indeed, condensates predicted to transiently accommodate a larger excess of RNA would also be expected to recruit additional positive counterions to maintain overall electroneutrality. The stable condensate with degradation exhibits an RNA-rich compositional imbalance, reaching a negative-to-positive charge ratio of approximately two. When degradation is absent, RNA polymerisation drives the condensate to even larger compositional imbalances, but only transiently before condensate dissolution. In the persistent condensate with degradation, its composition corresponds to an RNA/FUS mass ratio of ∼0.15, compared with the equilibrium optimum of ∼0.1 observed previously for FUS/RNA condensates^23^.

Lastly, our simulations with the REACT-RNA framework enable us to resolve the internal molecular architecture of evolving non-equilibrium condensates containing RNA chains of variable length, revealing an important contribution of interfacial properties to the resilience of condensates to excess negative charge. To understand how the balance between polymerization and degradation defines the steadystate spatial organisation of RNA molecules within the condensate, we computed the average RNA chain length as a function of the distance from the condensate centre for the simulation with RNA degradation enabled, considering only the behaviour at longer times (*t >* 500 ns). Figure 4(h) shows this quantity together with the density profile, facil-itating identification of the condensate shape and interface position. Despite RNA lengths being dynamically edited through ongoing polymerization and degradation, we find that they spontaneously adopt an inhomogeneous spatial organisation under non-equilibrium conditions. Specifically, the average RNA chain length decreases monotonically with increasing distance from the condensate center. This result aligns with previous computational predictions of equilibrium condensates^23,52^ and reflects a spatial segregation of RNA molecules by length, in which long RNA molecules are preferentially buried within the condensate core, while shorter RNA molecules accumulate near the interface. To our knowledge, this length-dependent spatial partitioning has not yet been directly demonstrated experimentally and therefore constitutes a clear, experimentally testable prediction of our model. Long RNA strands possess higher effective valency and therefore establish a larger number of favourable RNA–FUS interactions, increasing the density of molecular connections within the condensate and lowering the bulk enthalpy of the dense phase. In contrast, shorter RNA molecules act as effective surfactants^23^, preferentially localising at the interface where they reduce the interfacial free energy and surface tension. Accordingly, the steady-state inhomogeneous RNA length distribution simultaneously enhances condensate stability through a highly connected core and lowers the energetic cost of maintaining the condensate interface.

Biomolecular condensate interfaces are increasingly recognized as chemically distinct environments that can profoundly influence molecular transport, interactions, and reaction kinetics^30,53–56^. We therefore asked whether the condensate bulk and interface differ in their propensity to drive RNA polymerization in our simulations by computing the probability of RNA polymerization as a function of the distance from the condensate centre. In Figure 4(i) we show this probability alongside the all-particle density profile (continuous line) and the terminal RNA residues (also including free nucleotides)(dashed line). This probability exhibits a pronounced maximum at ∼100 Å from the condensate centre of mass, coinciding with the condensate interface, which is identified by the decay of the all-particle density profile (blue continuous line). This peak is accompanied by an enrichment of RNA precursors and terminal RNA residues (including free nucleotides; dashed line) relative to the condensate core. Together, this indicates that RNA synthesis occurs preferentially at the condensate interface.

The preferential localization of RNA polymerization at the interface arises from the combined effects of reactant supply, condensate molecular organization, and enhanced interfacial dynamics. In our simulations, free nucleotides are supplied from the dilute phase, and therefore they penetrate the condensate through the interface, generating a continuous reactive flux into this region. At the same time, shorter RNA chains preferentially localize at the interface because they induce a lower interfacial free-energy penalty^23^, enriching RNA strand termini where elongation occurs. In addition, molecules exhibit a characteristic enhanced molecular diffusion at condensate interfaces as recently shown by us^30^. Overall, these three effects create a highly reactive environment at the interface that promotes nucleotide incorporation and RNA chain growth. These findings constitute a further testable prediction of our model: the condensate interface functions as a reactive locus where nucleotide uptake, RNA elongation, and molecular exchange between the dense and dilute phases are simultaneously maximized, thereby explaining the interfacial enrichment of newly synthesized short RNAs (Fig. 4(h–i)). As RNA strands continue to elongate, they progressively migrate away from the interface toward the condensate core.

Together, these results show that the coupling between RNA polymerisation, degradation, and continuous nucleotide influx drives condensates into non-equilibrium steady states with RNA concentrations above the previously reported equilibrium optimum. By dynamically regulating RNA length, concentration, and interfacial composition, these coupled processes delay the onset of the high-RNA re-entrant dissolution regime and allow condensates to transiently sustain elevated RNA concentrations and an RNA-rich compositional imbalance

This mechanism provides a physical basis for how transcriptional condensates could accommodate bursts of RNA production. Rather than dissolving immediately upon RNA accumulation, condensates can transiently buffer elevated RNA concentration and excess negative charge, provided degradation controls unbounded RNA growth and maintains optimal RNA lengths. Dissolution therefore emerges when degradation can no longer compensate continued RNA accumulation and the associated loss of intermolecular connectivity.

### E. RNA Polymerization can Dissolve Aged FUS Condensates

In previous work, we developed a computational framework capable of capturing condensate ageing mediated by the formation of inter-protein β-sheets within low-complexity aromatic-rich kinked segments (LARKS)^31,32,57,58^. These interactions progressively transform liquid-like condensates into more solid- or gel-like assemblies with reduced molecular mobility^59,60^. Here, we use this framework to investigate how continuous RNA polymerisation remodels aged FUS condensates containing already formed inter-protein β-sheets (see SM section SVII for details).

To this end, we coupled REACT-RNA—connected to a nucleotide reservoir in the dilute phase—to a pre-formed and pre-aged FUS condensate containing inter-protein β-sheets distributed across the FUS low-complexity domain^61^. FUS aging simulations were performed below the critical solution temperature (T = 325 K), such that FUS first formed a condensate that subsequently aged through progressive accumulation of inter-protein β-sheets. Once ageing was established, RNA polymerisation via REACT-RNA was activated. As polymerisation proceeds, RNA molecules progressively accumulate within the condensate (Figure 5(a), middle), where they interact with FUS while simultaneously increasing RNA–RNA electrostatic repulsion. At longer timescales (*t >* 1500 ns; Figure 5(a), bottom), this repulsion destabilises the intermolecular connectivity of the aged condensate, which fragments into smaller clusters that remain partially stabilised by cross-β-sheet interactions. This behaviour is quantified by the monotonic decrease in the condensation degree parameter (Figure 5(b)) and by the emergence of long RNA chains (*>* 500 nucleotides; Figure 5(c)). Therefore, continuous RNA growth can destabilise even aged, gel-like condensates, revealing how non-equilibrium RNA polymerisation actively remodels kinetically arrested protein networks.

**Figure 5:**
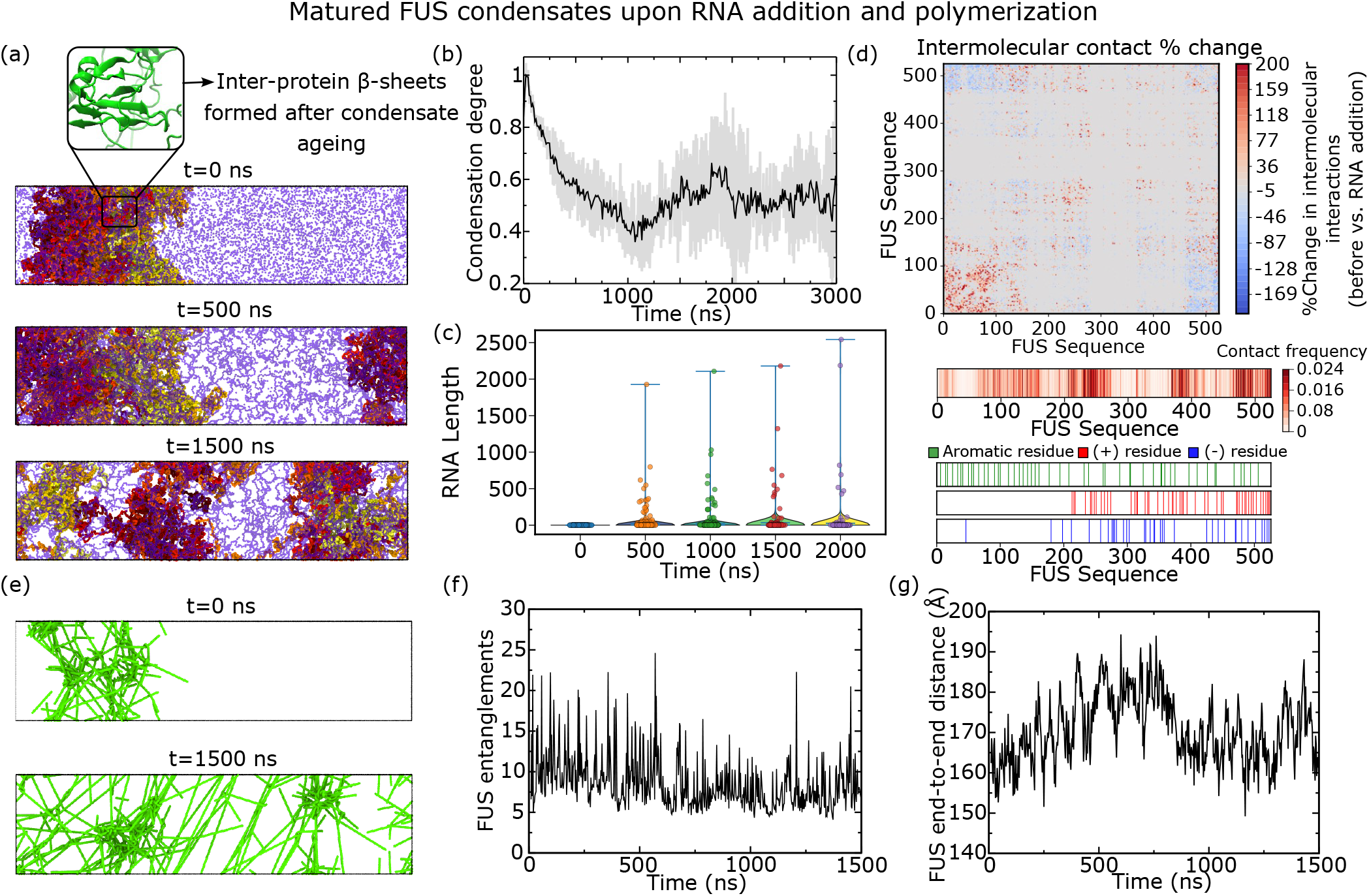
Aged FUS condensate dissolves in the presence of growing RNA. (a) Rendered images representing the time evolution of the aged FUS condensate (red tones) as polymerizing RNA (blue) emerges. (b) Normalized condensation degree parameter as a function of time. (c) Time-resolved RNA length distributions from violin plots (d) Top: Intermolecular contact map difference for FUS-FUS intermolecular interactions expressed in %. The initial and final steps are t=0 ns (aged condensate) and t=500 ns (upon RNA insertion). Bottom: FUS-RNA intermolecular contacts frequency map measured upon RNA growth (t=500 ns), along with FUS sequence residue map where the aromatic (green), positive (red) and negatively charged (blue) residues are highlighted. (e) Primitive path analysis showing the entanglements formed by the FUS/RNA condensate at times t = 0 ns and t = 1500 ns. For this analysis we have considered RNA strands only above 20 nucleotides of polymerization degree to avoid over-representation of effectively non-entangled strands (see Section SVIII for details). (f) Total number of entanglements formed by FUS as a function of time obtained from primitive path analysis. (g) Time dependence of FUS end-to-end distance.

To elucidate the molecular origin of this remodelling process, we analysed changes in intermolecular contacts between the initial aged condensate and the RNA-perturbed state at *t* = 500 ns (Figure 5(d)). Most FUS–FUS interactions remain relatively unchanged, except within the low-complexity domain, where intermolecular contacts increase. In contrast, interactions between the low-complexity domain and the C-terminal RGG3 region decrease, consistent with a redistribution of intermolecular interactions towards FUS– RNA binding. Analysis of the FUS–RNA contact map further shows that RNA preferentially binds positively charged regions enriched in arginine and lysine residues, particularly residues 240–270, 380–400, and 470–526. Significant RNA interactions are also observed within the low-complexity domain, likely promoted by its aromatic residue content. Thus, RNA growth redistributes intermolecular connectivity within the aged condensate while preserving residual interactions within LARKS-containing regions.

To characterise the topological changes to the network of intermolecular connections accompanying condensate remodelling, we performed primitive path analysis (PPA) along the simulation trajectory^62,63^ (see SM section SVIII for details). Rendered PPA configurations (Figure 5(e)) reveal that the entanglement network associated with inter-protein β-sheets partially persists even after condensate destabilisation, although the connectivity between different network nodes becomes markedly reduced. Consistently, residual FUS–RNA clusters remain stabilised by local inter-protein β-sheet interactions, while the total number of FUS entanglements remains approximately constant throughout the simulation (Figure 5(f)). In contrast, individual FUS conformations exhibit only modest changes, with the end-to-end distance varying by less than 10% during RNA accumulation (Figure 5(g)). These results indicate that RNA-driven dissolution primarily remodels intermolecular connectivity and network topology rather than inducing large conformational rearrangements of individual proteins.

These results reveal a non-equilibrium mechanism by which continuous RNA growth progressively weakens and remodels aged condensates through the coupled effects of RNA accumulation, multivalent protein–RNA interactions, and RNA–RNA electrostatic repulsion. Importantly, dissolution remains incomplete because local cross-β-sheet interactions preserve residual entangled clusters even after large-scale condensate destabilisation. Such electrostatically driven destabilisation mechanisms resemble previous observations for amyloid fibrils^64,65^ and suggest that RNA degradation may contribute to the remodelling or clearance of dysfunctional condensates in cellular environments. Given the established links between condensate ageing and neurodegenerative disease^66,67^, these findings provide a conceptual framework for understanding how non-equilibrium nucleic-acid fluxes may regulate the properties of pathological protein assemblies.

## III. CONCLUSIONS

In this work, we introduce REACT-RNA, a chemical-specific coarse-grained molecular dynamics framework that captures the dynamic formation of nucleotide–nucleotide bonds in protein-crowded environments. This approach enables direct simulation of RNA polymerisation, nucleotide exchange with a surrounding precursor reservoir, and RNA degradation within protein–RNA condensates. By coupling these reactions to sequence-specific protein–RNA interactions, REACT-RNA provides a molecular-resolution method for studying how RNA length, concentration, and degradation emerge dynamically within non-equilibrium condensates.

RNA polymerisation can promote condensate formation by generating multivalent RNA strands that bridge RNA-binding proteins and enhance intermolecular connectivity. Under limited nucleotide supply, this mechanism stabilises FUS condensates under near-critical conditions. In the pres-ence of a constant flux of nucleotides, RNA polymerisation drives transient MED1–RNA coacervation. These results are in agreement with experimental and computational studies identifying RNA as a key regulator of FUS and MED1 phase behaviour^11,17,19,24^. However, RNA-mediated condensation is inherently self-limiting. Continued RNA growth eventually saturates protein–RNA binding interactions and enhances the effective RNA–RNA repulsive interactions of the model, driving re-entrant dissolution of both FUS and MED1 condensates at high RNA concentrations, consistent with previous observations of RNA-driven condensate dissolution^16^. Because our model employs effective screened electrostatic interactions without explicit counterions, the actual range of protein/RNA concentration of this mechanism should be interpreted as model-dependent, given that it does not account for processes such as counterion condensation or ion-mediated RNA–RNA interactions that may influence the behaviour of highly RNA-rich condensates. Nevertheless, the underlying mechanism remains unchanged: RNA polymerisation acts as a dynamic control parameter that can both promote and destabilise condensates depending on the evolving RNA length and concentration.

A central result of this work is that RNA degradation and nucleotide flux reshape this re-entrant behaviour. When RNA polymerisation is sustained by a nucleotide reservoir, degradation counteracts RNA overaccumulation and dynamically selects RNA length distributions that preserve condensate connectivity. As a result, the RNA composition of condensates actively transcribing RNA is history dependent, reflecting the prior balance between RNA production, degradation, and molecular fluxes. In this scenario, condensates can remain stable while sustaining higher RNA concentrations and larger RNA-rich compositional imbalances (as quantified by the negative-to-positive charge ratio in our model) than expected from equilibrium condensates. This suggests that RNA degradation provides within our modelling framework a physical mechanism by which transcriptional condensates could buffer bursts of RNA production without immediately crossing into the high-RNA dissolution regime. Furthermore, it suggests that while RNA sequence identity is defined by the underlying transcript, the RNA length distribution within a transcriptional condensate could emerge as a dynamic non-equilibrium property regulated by the coupled balance between RNA polymerisation versus degradation rates, molecular exchange, and condensate partitioning.

A key prediction of our model is that RNA polymerisation occurs preferentially at the condensate interface. This interfacial enrichment of RNA polymerisation arises from the combination of three distinct effects: continuous influx of free nucleotides entering the condensate from the dilute phase, enrichment of reactive RNA chain termini near the interface, and the enhanced molecular mobility characteristic of condensate interfaces^30^. Together these effects generate a reactive region in the condensate in which nucleotide uptake, RNA chain elongation, and molecular exchange between the dense and dilute phases are simultaneously maximised. These findings provide a molecular and thermo-dynamic explanation for the experimental observation that transcription occurs at or near the interface of transcriptional condensates^11,13,14^. This result also exemplifies how as RNA polymerisation reshapes condensates, condensates in turn promote the reactive molecular configurations that favour RNA polymerisation.

The localisation of RNA polymerisation at the interface is accompanied by a spatially organised RNA population. The resulting steady-state condensates exhibit an inhomogeneous RNA length organisation, in which shorter, newly synthesised RNAs accumulate near the interface, whereas longer RNAs preferentially partition towards the condensate core. This spatial organisation provides a mechanism by which condensates can simultaneously regulate their interfacial properties—through short RNAs that act as surfactants and lower the interfacial free energy—and maintain a highly connected internal molecular network sustained by long, high-valency RNAs. Although this length-dependent spatial partitioning remains to be directly tested experimentally, it constitutes a clear and experimentally falsifiable prediction of our model.

These results are consistent with growing evidence that condensate interfaces possess distinct molecular organisation and can actively regulate transport, molecular exchange, biochemical reactions and structural transitions^13,14,30,55^. In this context, our results suggest that transcriptional condensates may physically regulate RNA length distributions through the coupled balance between RNA polymerisation, degradation, and interfacial molecular fluxes.

We also show that continuous RNA growth can remodel aged, gel-like FUS condensates stabilised by inter-protein β-sheet interactions. In this regime, RNA polymerisation acts as a non-equilibrium perturbation that progressively injects negative charge and protein–RNA interactions into a kinetically arrested network, weakening intermolecular connectivity and fragmenting the aged condensate into smaller residual clusters. These results suggest that RNA dynamics may not only regulate the formation and dissolution of liquid-like condensates, but may also contribute to the remodelling of more arrested condensate states associated with cellular stress and neurodegenerative diseases^68^. In this context, our model predicts that charged biomolecules that are mutually self-repulsive yet interact favourably with aggregation-prone proteins could promote the destabilisation of pathological assemblies at sufficiently high concentrations. This prediction is consistent with previously proposed electrostatic mechanisms for amyloid fibril destabilisation^58,64,65,69^ but should be regarded as a speculative implication of the present model rather than as evidence that RNA-mediated remodelling or clearance of dysfunctional condensates occurs in cells. Furthermore, recent work has shown that condensate interfaces can favour structural transitions such as fibril formation and ageing^30,53,55,57^, reinforcing the idea that interfaces act as active regulatory regions controlling both condensate chemistry and material-state transitions.

An important aspect of the REACT-RNA framework is that RNA polymerization is controlled by several modelling parameters, including the polymerization distance cutoff, the protein-contact criterion for polymerization, the reservoir concentration, the RNA degradation probability, and the overall system density. These variables collectively determine the effective polymerization rate and therefore influence the balance between RNA production and degradation. Rather than representing arbitrary tuning parameters, they were selected with two goals: first, to fit experimentally observed qualitative conditions (e.g., solution concentration of both proteins and nucleotides) while allowing RNA growth, condensate formation, and dissolution to be characterized within computationally accessible simulation times; and second, to ensure that each parameter has a physically motivated basis (i.e., relative molecular diffusion/polymerization rates), as detailed and justified in the Supplementary Material sections SXI and SXIII. Consequently, our conclusions should be interpreted within the framework of this protein-coupled reaction model. Within this framework, the observed feedback between RNA production and condensate organization, including the emergence of an optimal RNA regime, arises dynamically from the interplay between polymerization, degradation, and RNA-driven phase separation rather than from the imposition of an specific target RNA concentration or stoichiometry.

Overall, our work shows that RNA–protein condensates are governed by the coupled dynamics of RNA production, degradation, molecular fluxes, and interfacial organisation. RNA polymerisation injects mass, charge, and multivalency into condensates, whereas degradation limits overgrowth and regulates the RNA length distribution. Their competition defines the window over which condensates remain stable, delays the onset of re-entrant dissolution, and generates spatially organised non-equilibrium steady states. These findings provide a physical framework for understanding how transcriptional condensates may use RNA degradation and interfacial reactions to regulate their composition, charge state, internal architecture, and stability.

## Supporting information

Suppementary Material

## IV. DATA AVAILABILITY

The data that supports the findings of this study are available within the article and its Supplementary Material.

## V. CODE AVAILABILITY

LAMMPS scripts and initial configurations for all the simulations described throughout this manuscript can be found in our repository https://doi.org/10.5281/zenodo.19585081

## VI. ACKNOWLEDGEMENTS

S.-B. acknowledges funding from UKRI EPSRC under the UK Government’s guarantee scheme (EP/Z002028/1), following successful evaluation by the ERC (Consolidator Grant awarded to R.C.G.) under the European Union’s Horizon Europe research and innovation programme. A. R. T. acknowledges funding from the European Union Horizon 2020 research and innovation programme (grant agreement 803326 to R.C.-G.). J.R.E. acknowledges funding from the Ramon y Cajal fellowship (RYC2021-030937-I), the Spanish National Agency for Research (PID2022-136919NA-C33 and PID2025-169417NBC21), and the European Research Council (ERC) under the European Union’s Horizon Europe research and innovation program (grant agreement no. 101160499). J.R.E also acknowledges the CRIS Cancer Foundation for the research grant CRIS-CANCER-4332687 and the BBVA foundation (G999088J). Research in the Collepardo-Guevara lab is supported by the UK Research Innovation (UKRI) Engineering and Physical Sciences Research Council (EPSRC) [EP/Z002028/1]. This work has been performed using resources provided by Archer2 (https://www.archer2.ac.uk/) funded by EPSRC Tier-2 capital grant EP/P020259/e829. The authors also thankfully acknowledge RES computational resources provided by Mare Nostrum 5 through the activity 2024-3-0001.

## VIII. SUPPLEMENTARY MATERIAL

Additional simulation details, materials, and methods. Simulation algorithms detailed description, protein sequences and additional sections validating the variable parameters choice.

