## Supplementary material for "RNA synthesis and degradation regulate biomolecular condensates through non-equilibrium feedback": Suppementary Material

### SI. MPIPI-RECHARGED FORCE FIELD

We use the Mpipi-Recharged model, a residue-level coarse-grained force field for both protein and protein/RNA condensates [1]. In this model, each amino acid or nucleotide is represented by a single bead connected to adjacent residues by harmonic bonds. Globular domains of the proteins are treated as rigid bodies whose beads are fixed at the $C_\alpha$ of the corresponding Protein Data Bank (PDB). The potential energy is computed as the sum of pairwise bonded ($E_{\text{bonded}}$) and non-bonded ($E_{\text{non-bonded}}$) interactions as:

$$E = E_{\text{bonded}} + E_{\text{non-bonded}}. \tag{S1}$$

The intrinsically disordered regions (IDRs) are modelled as fully flexible polymers and these are connected to the globular domains. The bonded potential is written as:

$$E_{\text{bonded}}(r_{ij}) = \sum_{ij} k(r_{ij} - r_0)^2, \tag{S2}$$

where $r_{ij}$ is the distance between the connected beads, $r_0 = 3.81\,\text{Å}$ and $r_0 = 5.00\,\text{Å}$ are the equilibrium bond lengths for protein and RNA, respectively. The spring constant $k = 9.60\,\text{kcal}\cdot\text{mol}^{-1}\cdot\text{Å}^{-2}$. The sum runs over all paired residues.

Non-bonded interactions consist of the sum of the hydrophobic interaction and electrostatic interaction. The hydrophobic interaction is given by the Wang–Frenkel (WF) potential [2] that accounts for short-ranged excluded-volume repulsion and long-ranged attraction. This potential is defined as

$$E_{\mathsf{WF}}(r_{i,j}) = \sum_{ij} \epsilon_{ij}\alpha_{ij} \left[\left(\frac{\sigma_{ij}}{r_{ij}}\right)^{2\mu_{ij}} - 1\right] \left[\left(\frac{R_{ij}}{r_{ij}}\right)^{2\mu_{ij}} - 1\right]^{2\nu_{ij}}, \tag{S3}$$

where

$$\alpha_{ij} = 2\nu_{ij}\left(\frac{R_{ij}}{\sigma_{ij}}\right)^{2\mu_{ij}} \left\{\frac{2\nu_{ij}+1}{2\nu_{ij}\left[\left(\frac{R_{ij}}{\sigma_{ij}}\right)^{2\mu_{ij}} - 1\right]}\right\}^{2\nu_{ij}+1}. \tag{S4}$$

Here $\sigma_{ij}$ is the pair-of-beads diameter, defined from the individual diameter ($\sigma_i$ and $\sigma_j$) assuming the Lorentz-Berthelot mixing rules (i.e., $\sigma_{ij} = (\sigma_i + \sigma_j)/2$). $R_{ij} = 3\sigma_{ij}$ is the cut-off distance for the $ij$-th interaction. The interaction parameter $\epsilon_{ij}$ is defined for each specific amino acid pair based on our atomistic Potential of Mean Force calculations and bioinformatics data [1]. The exponent $\nu_{ij}$ is set to 1 for all the pairs and $\mu_{ij}$ depends on the specific pair, ranging from 2 to 12 (see Ref. [1]). Notice that higher values of $\mu_{ij}$ lead to a steeper increase in the repulsive part of the potential. The interaction involving globular domains are reduced to account for the 'buried' interactions. In particular the interaction between flexible regions and globular domains are screened by a factor of $\sqrt{0.7}$ and the WF interaction between residues in globular regions is scaled down a factor of $0.7$.

The electrostatic interactions are described by the Yukawa potential [3] instead of the Debye–Hückel potential [4] originally employed in the Mpipi model. Avoiding the use of explicit charge values, this potential allows to modulate independently the strength of electrostatic interactions in a pair-specific basis. This potential is defined as

$$E_{\mathsf{electrostatic}} = \sum_{ij} \frac{A_{ij}}{r_{ij}} \exp(-\kappa r_{ij}), \qquad r_{ij} < r_c, \tag{S5}$$

where $A_{ij}$ is the interaction parameter, $\kappa$ is the screening due to ions, and $r_{ij}$ is the distance between interacting pairs. The cut-off for the electrostatic interaction, $r_c$, is set to 3.5 nm. As recently shown by Tesei and Lindorff-Larsen [5], cutoff values above 3 nm results in accurate predictions for a very similar force field, preserving the measured properties including the saturation concentration and the radius of gyration. The screening parameter $\kappa$ is expressed in a explicit way as $\kappa = \sqrt{8\pi B c_s}$, where $c_s$ is the salt concentration—normally set to 150 mM of NaCl—and $B = e_0^2/4\pi k_B T \varepsilon_0 \varepsilon_r$ is the Bjerrum length. The relative dielectric constant $\varepsilon_r$ varies with temperature according to the empiric formula [6]:

$$\varepsilon_r(T) = \frac{5321}{T} + 233.760 - 0.9297T + 1.417 \cdot 10^{-3} T^2 - 8.292 \cdot 10^{-7} T^3, \tag{S6}$$

for $T$ in Kelvin. The pair-specific optimized values of $A_{ij}$ of the Yukawa potential can be found in Ref. [1]. In the Mpipi-Recharged model, such parameters indicate that the interaction between oppositely charged pairs is significantly stronger than those between identically charged pairs.

### SII. SIMULATION DETAILS

All simulations were conducted using the LAMMPS software [7]. All our calculations were performed in the NVT ensemble. For the simulations in which the RNA reservoir (see section SV for details) and RNA degradation (see section SVI for details), short runs in the NVT are performed, followed by a potential change in the total number of particles, therefore not conserving this quantity over time. For simulations with no rigid bodies (no globular domains such as MED1 LCD), we employing a Nosé-Hoover thermostat [8] with a relaxation time of 100 fs. When using rigid bodies, such as the case of FUS (see below) Langevin thermostat [9] with a relaxation time of 5 ps was used. The time step for the Verlet integration is 10 fs. We fix the electrolyte solution ionic strength to 150 mM for FUS simulations and 75mM for MED1, in agreement with the experiments by Henninger *et al.* that highlight the salt effect in MED1/RNA condensation [10]. FUS simulations at the critical temperature (Figures 2 and 4 in the main text) are performed at 337 K, while the ones shown for MED1 are performed at 250 K. The temperature choice for MED1/RNA was set so that we would observe condensation over a computationally accessible timescale, since we did not observe condensation at temperatures above. Nonetheless, this ~20 K discrepancy in phase separation temperature with respect to the experimental conditions is within the expected difference observe for residue-resolution coarse-grained models [11]. Additionally, aged FUS condensate simulations (Figure 5) are performed at T = 325 K, below $T_c$ so that an aged condensate is first formed in absence of RNA. Our simulations contain a changing amount of particles, and we have consequently verified that the global system density remains between 0.08 and 0.16 $g{\cdot}cm^{-3}$, in order to observe phase separation, according to the phase diagram from Figure 2a.

When simulating FUS, we keep the globular domains constrained using the rigid bodies integrator for those segments. The following Protein Data Bank (PDB) codes were used to build the globular structured domains of FUS (residues from 285–371 (PDB code: 2LCW) and from 422–453 (PDB code: 6G99)).

Below are the sequences of both FUS and MED1 LCD

FUS sequence:

| | | | | | | |
|---|---|---|---|---|---|---|
| 1 | 11 | 21 | 31 | 41 | 51 | 61 |
| MASNDYTQQA | TQSYGAYPTQ | PGQGYSQQSS | QPYGQQSYSG | YSQSTDTSGY | GQSSYSSYGQ | SQNTGYGTQS |
| 71 | 81 | 91 | 101 | 111 | 121 | 131 |
| TPQGYGSTGG | YGSSQSSQSS | YGQQSSYPGY | GQQPAPSSTS | GSYGSSSQSS | SYGQPQSGSY | SQQPSYGGQQ |
| 141 | 151 | 161 | 171 | 181 | 191 | 201 |
| QSYGQQQSYN | PPQGYGQQNQ | YNSSSGGGGG | GGGGGNYGQD | QSSMSSGGGS | GGGYGNQDQS | GGGGSGGYGQ |
| 211 | 221 | 231 | 241 | 251 | 261 | 271 |
| QDRGGRGRGG | SGGGGGGGGG | GYNRSSGGYE | PRGRGGGRGG | RGGMGGSDRG | GFNKFGGPRD | QGSRHDSEQD |
| 281 | 291 | 301 | 311 | 321 | 331 | 341 |
| NSDNNTIFVQ | GLGENVTIES | VADYFKQIGI | IKTNKKTGQP | MINLYTDRET | GKLKGEATVS | FDDPPSAKAA |
| 351 | 361 | 371 | 381 | 391 | 401 | 411 |
| IDWFDGKEFS | GNPIKVSFAT | RRADFNRGGG | NGRGGRGRGG | PMGRGGYGGG | GSGGGGRGGF | PSGGGGGGGQ |
| 421 | 431 | 441 | 451 | 461 | 471 | 481 |
| QRAGDWKCPN | PTCENMNFSW | RNECNQCKAP | KPDGPGGGPG | GSHMGGNYGD | DRRGGRGGYD | RGGYRGRGGD |
| 491 | 501 | 511 | 521 | | | |
| RGGFRGGRGG | GDRGGFGPGK | MDSRGEHRQD | RRERPY | | | |

MED1 LCD sequence:

| | | | | | | |
|---|---|---|---|---|---|---|
| 1 | 11 | 21 | 31 | 41 | 51 | 61 |
| EHHSGSQGPL | LTTGDLGKEK | TQKRVKEGNG | TSNSTLSGPG | LDSKPGKRSR | TPSNDGKSKD | KPPKRKKADT |
| 71 | 81 | 91 | 101 | 111 | 121 | 131 |
| EGKSPSHSSS | NRPFTPPTST | GGSKSPGSAG | RSQTPPGVAT | PPIPKITIQI | PKGTVMVGKP | SSHSQYTSSG |
| 141 | 151 | 161 | 171 | 181 | 191 | 201 |
| SVSSSGSKSH | HSHSSSSSSS | ASTSGKMKSS | KSEGSSSSKL | SSSMYSSQGS | SGSSQSKNSS | QSGGKPGSSP |
| 211 | 221 | 231 | 241 | 251 | 261 | 271 |
| ITKHGLSSGS | SSTKMKPQGK | PSSLMNPSLS | KPNISPSHSR | PPGGSDKLAS | PMKPVPGTPP | SSKAKSPISS |
| 281 | 291 | 301 | 311 | 321 | 331 | 341 |
| GSGGSHMSGT | SSSSGMKSSS | GLGSSGSLSQ | KTPPSSNSCT | ASSSSFSSSG | SSMSSSQNQH | GSSKGKSPSR |
| 351 | 361 | 371 | 381 | 391 | 401 | 411 |
| NKKPSLTAVI | DKLKHGVVTS | GPGGEDPLDG | QMGVSTNSSS | HPMSSKHNMS | GGEFQGKREK | SDKDKSKVST |
| 421 | 431 | 441 | 451 | 461 | 471 | 481 |
| SGSSVDSSKK | TSESKNVGST | GVAKIIISKH | DGGSPSIKAK | VTLQKPGESS | GEGLRPQMAS | SKNYGSPLIS |
| 491 | 501 | 511 | 521 | 531 | 541 | 551 |
| GSTPKHERGS | PSHSKSPAYT | PQNLDSESES | GSSIAEKSYQ | NSPSSDDGIR | PLPEYSTEKH | KKHKKEKKKV |
| 561 | 571 | 581 | 591 | 601 | 611 | 621 |
| KDKDRDRDRD | KDRDKKKSHS | IKPESWSKSP | ISSDQSLSMT | SNTILSADRP | SRLSPDFMIG | EEDDDL |

### SIII. DIRECT COEXISTENCE METHOD AND PHASE DIAGRAM DETERMINATION

We compute the phase diagram and critical temperature for phase separation for the FUS through NVT simulations via the Direct Coexistence method, where both condensed and diluted phase coexist under the equilibrium conditions. We perform simulations below the critical temperature, and measure the density of the coexisting condensed and diluted phases. When the phase diagram is calculated via this method, the critical point of the phase diagrams is estimated using the universal scaling law of coexistence densities near a critical point [12]

$$\big(\rho_l(T) - \rho_v(T)\big)^{3.06} = d\left(1 - \frac{T}{T_c}\right) , \tag{S7}$$

and the law of rectilinear diameters [13]

$$\frac{\rho_l(T) + \rho_v(T)}{2} = \rho_c + s_2(T_c - T) , \tag{S8}$$

where $\rho_l$ and $\rho_v$ refer to the coexisting densities of the condensed and diluted phases respectively, $\rho_c$ is the critical density, $T_c$ is the critical temperature, and $d$ and $s_2$ are fitting parameters.

### SIV. RNA POLYMERIZATION ALGORITHM

In the main text we describe the algorithm that allows the dynamic formation of nucleotide-nucleotide bonds throughout the simulation. More specifically, single nucleotides and terminal beads of RNA chains are capable of doing so. For this purpose, we use the *fix bond/react* command in LAMMPS. As a coordination criteria, nucleotides must be within a 7.5 Å in order for the bond to be created. Moreover, both nucleotides must also be within a 10 Å cutoff distance from any amino acid residue, in order to mimic the condensate-driven RNA elongation. We build the necessary template and map files required by LAMMPS for the bond creation process to occur. Moreover, we also implement a second reaction that remaps the molecule ID from the merging molecules so that the molecule ID is updated to be a single one of the two. Example files of this algorithm can be found in our repository https://doi.org/10.5281/zenodo.19585081 .

### SV. RNA INSERTION/DELETION ALGORITHM

In the main text we show simulations in which the concentration of RNA is kept constant in the dilute phase. This concentration-maintenance protocol does not conserve the number of particles in the simulation over time. The analysis is performed external to LAMMPS. We perform consecutive 5 ns simulations in LAMMPS using the $jump$ command. After one loop is finished, we run a external code which: (1) Computes the center of mass of the simulation (accounting for the presence of periodic boundary conditions in the 3 directions) (2) Evaluates the density profile (dividing the box into 100 slabs) from the center of mass in the elongated direction. The density profile is averaged in both directions. The diluted phase bound is set at the point at which the density profile reaches 15% of the value at the center. Therefore, the diluted phase is defined as the region further than such distance from the center of mass. (3) Then the algorithm calculates whether there is excess or deficiency of RNA with respect to a given reference value, which we set to 0.0001 nucleotides/Å$^3$. (4) We perform random deletion of RNA molecules in which all the nucleotides are contained in the diluted phase in case we find excess of RNA, until we reach the reference concentration. Either single nucleotides or RNA chains containing multiple nucleotides can be deleted as long as every residue is part of the diluted phase. While RNA chains can be deleted in this procedure, only single nucleotides are inserted. If we find deficiency, we insert single nucleotides at random positions within the diluted phase until reaching the target concentration. In this way we simulate the condensate in contact with a RNA nucleotides reservoir. Differently to the RNA degradation routine described in the following section, this procedure is intended to simulate the coexistence of a condensate with a RNA crowded environment only, while the RNA degradation algorithm is capable of targeting all RNA chains present in the system (including those inside the condensate).

### SVI. RNA DEGRADATION AT DIFFERENT PROBABILITIES

In Results section D of the main text we introduce the algorithm that deletes RNA molecules given a probability, which is given as an input parameter. More specifically, we use the following commands in our LAMMPS script that perform this task:

```
group rnastatic type 44 55 56
delete_atoms random fraction 0.001 yes rnastatic NULL 12345 bond yes mol yes
```

Here 0.001 is the 0.1% probability of deleting atoms from the RNA particles group defined in the first line. Therefore, particles are selected with such probability and afterwards, the entire molecules to which these particles belong are removed. Since our LAMMPS script is read in a 5 ns loop (see files in repository https://doi.org/10.5281/zenodo.19585081), this command is performed every 5 ns.

Moreover, we performed further simulations in which we sample different values of this probability, which account for different scenarios in which RNA molecules are degraded at different rates. No RNA degradation leads to completing the RNA transcription feedback loop, where in excess of RNA, phase separation is inhibited. We tested 5 different RNA degradation probabilities ranging from $10^{-1}$ to $10^{-6}$, and found that low deletion probabilities can allow sufficient RNA growth to lead to condensate dissolution mediated by RNA excess, while high ones keep the condensate stable over time (Figure S1).

### SVII. AGEING ALGORITHM

We perform our ageing simulations allowing the formation of inter-protein $\beta$-sheets to take place, according to the scheme developed previously [14–16]. We first identified the regions of the FUS sequence that are prone to forming inter-protein

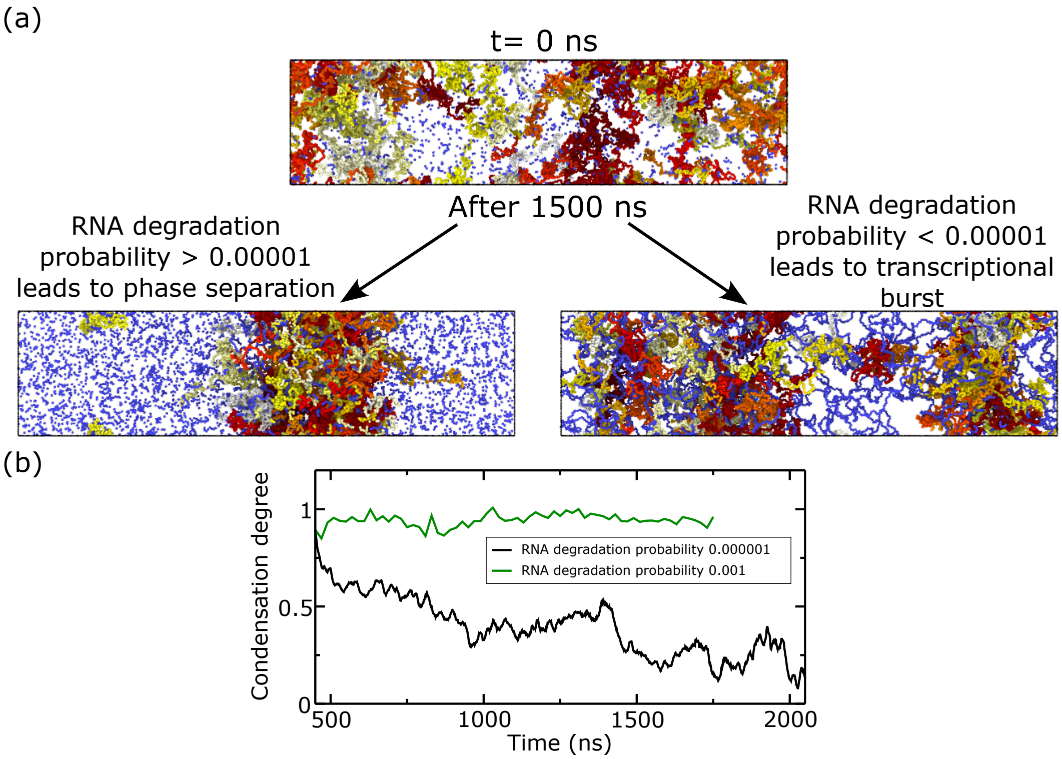


Figure S1: (a) Rendered images depicting the consequence of imposing different RNA degradation probabilities into our algorithm after simulating 1500 ns. (b) Condensation degree as a function of time for different cases above and beyond the degradation probability allowing stable condensation over time, as indicated in the label.

secondary structures, as reported in [17]. These are low-complexity aromatic-rich kinked segments (LARKS), and in the FUS LCD sequence we can find the following: $_{37}$SYSGYS$_{42}$ (PDB code 6BWZ), $_{54}$SYSSYGQS$_{61}$ (PDB code 6BXV), and $_{77}$STGGYG$_{82}$ (PDB code 6BZP) [17]. In Ref. [14] the potential of mean force of the structured and disordered conformations of such regions was calculated, indicating us what the interaction difference between before and after such structure is formed is. Our ageing simulations consist of non-equilibrium Molecular Dynamics runs in which, when the aforementioned regions are found in space, an effective change in the pairwise interaction of the involved residues (according to the PMF calculations) is performed. We use a distance criterion, in which at least the central residues of four LARKS of the same kind must meet within a cutoff distance of 14 Å. This distance is chosen so that the formation of the inter-protein $\beta$-sheets takes place within an accessible timescale. This distance is shorter but close to the Wang-Frenkel potential cutoff used for the non-charged residue-residue interactions. Nonetheless this choice is justified due to the fact that we measure the coordination of the central residue of the LARKS only, meaning that when this criterion is met, multiple residues of the LARKS belonging to different protein replicas are in close contact. Moreover, the requirement of at least four LARKS of different protein replicas meeting simultaneously is set so that the structures reported in Ref. [17] can be sustained. In this way we model the formation of inter-protein $\beta$-sheets. This is done with the *fix bond/react* command [18] available in LAMMPS. This algorithm has been previously used by us to describe the ageing process of FUS [16, 19–21], and example files of the script and input files for FUS ageing can be found in the repository https://doi.org/10.5281/zenodo.15168500

### SVIII. PRIMITIVE PATH ANALYSIS

We analyze the network connectivity of the FUS-RNA systems after maturation using the NVT trajectories after ageing (see Section SVII) and the primitive path analysis (PPA) method [22], modified in Tejedor *et al.* [16]. In order to keep the entanglements in a reasonable regime we delete all the RNA chains below 20 nucleotides long. Then, the PPA algorithm minimizes the contour length of all chains in the system while keeping the terminal residues of the monomers fixed and, in this way, preserving the network topology by preventing chains from crossing each other. The energy minimization is performed with a tolerance of $10^{-7}$ for both the energy and the force, allowing up to $5 \times 10^5$ iterations. After minimization, we obtain from each trajectory configuration several structural topological observables considering chains with lengths

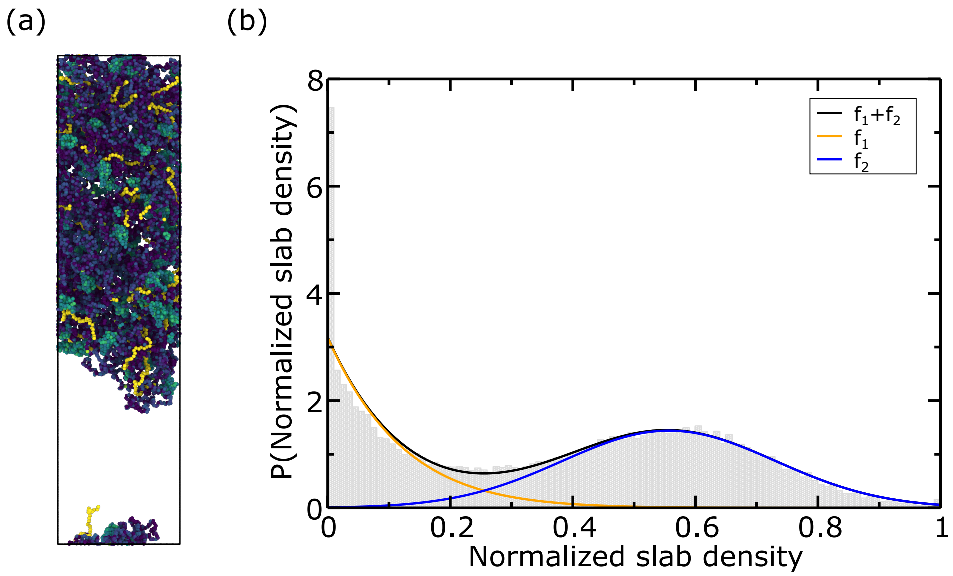


Figure S2: (a) Representative snapshot of the DC simulation analyzed to determine the optimal threshold for the condensation degree parameter. (b) Normalized slab density histogram. The black curve indicates the fit from the histogram data to $f_1+f_2$, while the orange and blue curves indicate the individual functions.

within the specified range (RNA longer or equal 20 nucleotides). In particular, we calculate the mean squared end-to-end distance $\langle \mathbf{R}^2 \rangle$ and the primitive path contour length $L_{pp}$ are directly calculated of the resulting configuration. The tube diameter $a$ is defined through the relation $\langle \mathbf{R}^2 \rangle = a \cdot L_{pp}$ [23]. The average number of entanglements per chain is then estimated as $Z = L_{pp}/a$.

### SIX. OPTIMIZING NORMALIZED DENSITY THRESHOLD FOR THE CONDENSATION DEGREE PARAMETER

In order to determine which density threshold to use in our simulations to determine the progress of the condensation or dissolution process, we analyse a DC trajectory (representative rendered image shown in Figure S2) and calculate the histogram of the normalized density found for each slab for all the independent configurations. We used the simulation described in Figure 2 of the main text at t $>$ 1000 ns, where we clearly have a condensed-dilute coexistence. Assuming that the normalized density is distributed normally, we perform a non-linear fit to a sum of normal distribution functions $f_1+f_2$, where

$$f_1 = \frac{1}{\sqrt{2\pi\sigma_1^2}} exp\left(-\frac{(x-\mu_1)^2}{2\sigma_1}\right) \text{ , } and \text{ } f_2 = \frac{1}{\sqrt{2\pi\sigma_2^2}} exp\left(-\frac{(x-\mu_2)^2}{2\sigma_2}\right). \quad \text{(S9)}$$

We have $\sigma_1, \sigma_2, \mu_1$ and $\mu_2$ as free parameters in the non-linear fit. We obtain that $f_1$ and $f_2$ cross at x=0.256, therefore such value minimizes the amount of slabs incorrectly labelled as condensed or diluted. Consequently, we use this value in the condensation degree parameter described in the main text.

### SX. CONTACT MAPS DEPENDENCE ON THE CUTOFF

The contact maps shown in the main text were calculated using 1.2 $\sigma_{ij}$ as the cutoff distance to consider that two residues have an effective interaction. Here we show the same contact maps analyzed with different cutoff, and show that despite the absolute frequencies variation, the qualitative conclusions drawn from all of them remain valid.

In Figure S3 we show the intermolecular MED1-RNA contact map computed for threshold values ranging between 1.1 to 1.5 $\sigma_{ij}$. As it can be seen from this representation, the regions more frequently establishing contacts remain almost identical, with the total frequency increasing as the cutoff distance increases.

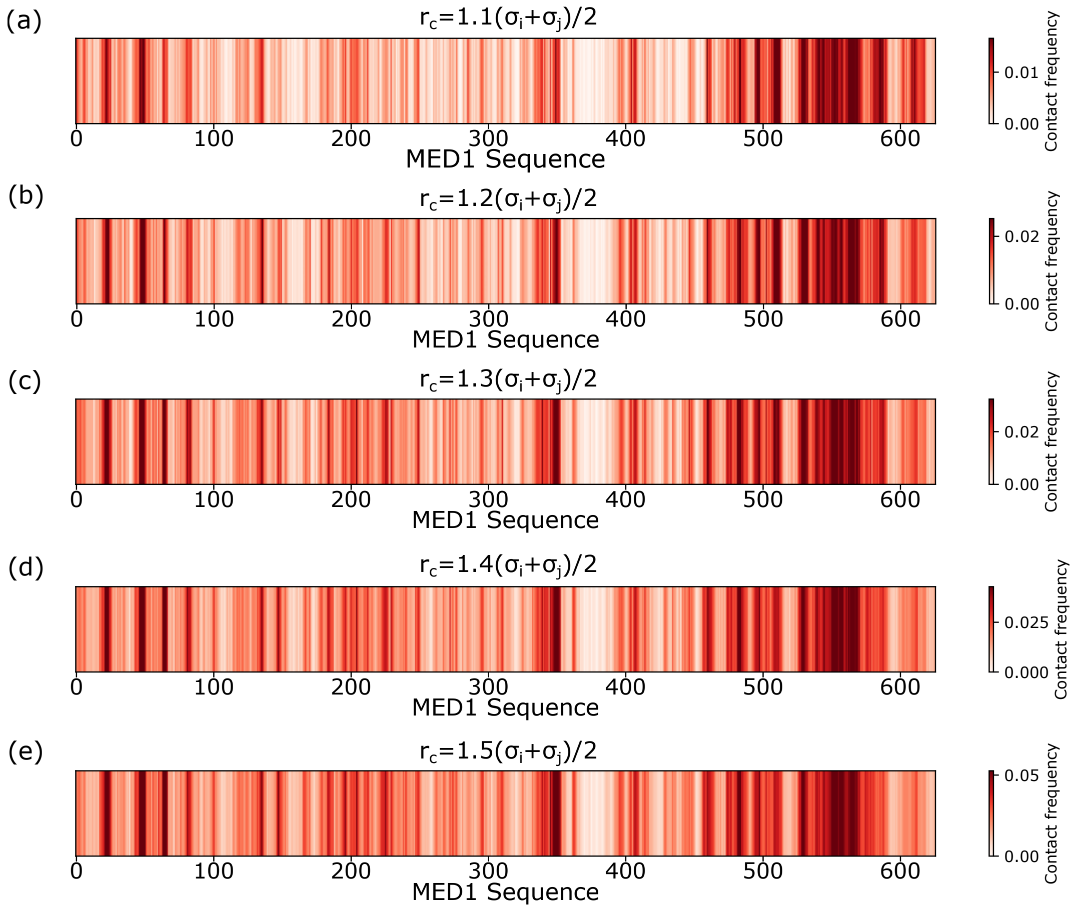


Figure S3: (a)-(e) MED1–RNA intermolecular contacts frequency along the MED1 sequence, where the cutoff distance to consider two residues as neighbours ($r_c$) is indicated on top of the corresponding contact map.

### SXI. VARIATIONS TO THE NUCLEOTIDE CONCENTRATION IN THE RESERVOIR

In this section we simulate a FUS/RNA system with no RNA degradation and a RNA reservoir concentration paired to experimentally reported in vivo [24]. In our simulations presented in the main text we set a RNA concentration in the diluted phase of 0.0001 nucleotides/Å$^3$, which is 166 mM. Here we pair the diluted phase to be of the experimentally observed in cells order of magnitude $\mathcal{O}$(5 mM) [24], imposing a diluted phase RNA concentration of 0.000005 nucleotides/Å$^3$ (8.3 mM).

In Figures 4(a)-(c) of the main text we observed and characterized the condensation of FUS mediated by RNA growth, followed by its dissolution, a feedback loop being completed in 1000 ns of simulation time. In Figure S4 we show the time evolution of the condensation degree as well as the longest RNA chain length present in the simulation. The evolution of this system is identical to that of Figure 3(a)-(c) (main text), however it requires over 5 times of simulation time. This result proves that, qualitatively and with enhanced dynamics, our choice of the RNA concentration surrounding the condensate is capable of recapitulating the behaviour appearing under conditions of physiological relevance. We further note that experimental studies have also employed strategies to locally enhance RNA concentrations around transcriptional condensates. For example, Henninger *et al.* [10] engineered an RNA super-enhancer system to promote local RNA production in the vicinity of condensates, thereby increasing the concentration of nascent RNA surrounding the protein-rich phase. Although achieved through a different mechanism than our imposed RNA reservoir, this experimental approach similarly creates an RNA-enriched environment around the condensate, supporting the relevance of considering elevated local RNA availability when investigating RNA-driven condensate dynamics.

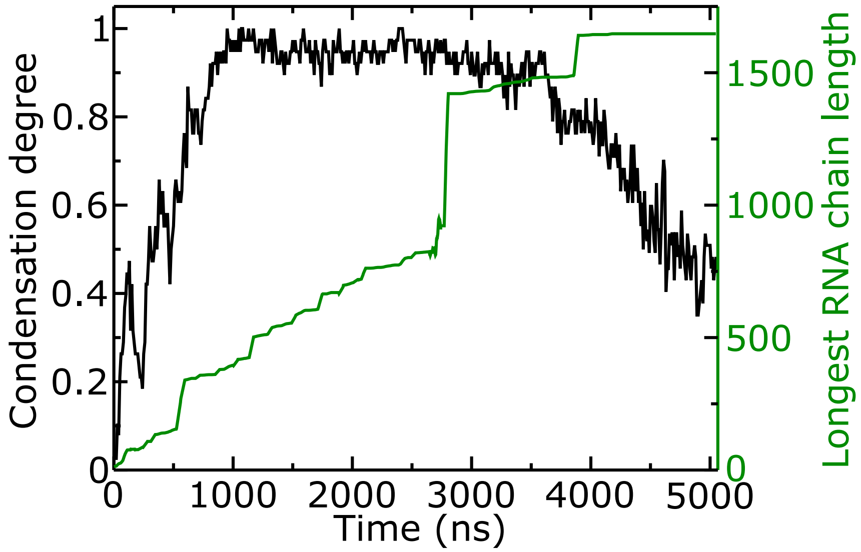


Figure S4: Condensation degree and longest RNA chain length present in the simulation as a function of time. Simulation carried out at the same conditions (337 K, no RNA degradation) as that shown in Figure 3(a)-(c) of the main text, varying the reference RNA concentration in the dilute phase to 0.000005 nucleotides/Å$^3$

### SXII. FUS CONDENSATION AT THE CRITICAL TEMPERATURE IS SUSTAINED BY RNA GROWTH

In Results section B of the main text we prove that RNA polymerization drives FUS condensation under critical temperature conditions. Here we provide further characterization of this process, as well as demonstrate the validity of this statement by providing comparision with a control scenario in which RNA polymerization is disabled, and all the nucleotides remain present in the simulation box. In Figure S5(a), we provide rendered images that clearly manifest that RNA polymerization is crucial for condensation under such conditions, as free nucleotides are not capable to bind FUS molecules alone. All the rendered images are accompanied by a density profile, where it can be seen that unless RNA lengthening drives condensation, these profiles remain flat, indicating an homogeneous distribution of the molecules along the simulation box, characteristic of a single dispersed phase. In Figure S5(b), we show the time evolution of the condensation degree described in the main text, computed for both the case with RNA polymerization enabled and disabled. In agreement with what we observe in panel (a), the condensation degree increases over time where RNA is allowed to grow, and remains flat at lower values for the scenario with free nucleotides. Lastly, in Figure S5(c), we plot the difference in density between the slabs with greater and lower density found in the simulation box ($\Delta\rho$) as a function of time. Also in agreement with panels (a) and (b), in the simulation with RNA polymerization disabled we find a flat curve that fluctuates over values below 0.2 g·cm$^{-3}$, which are characteristic of fluctuations found in a homogeneous system. Oppositely, the simulation in which RNA grows shows grater $\Delta\rho$ values above 0.3 g·cm$^{-3}$, indicating that this difference comes from the density difference between condensed and diluted phases.

### SXIII. RNA POLYMERIZATION CUTOFF CHOICE

In this section we aim to establish an optimal value for the cutoff distance required for two RNA residues (either terminal ones or single nucleotides) to form a new bond. We focus on the simplest scenario studied, in which no particle reservoir exchanges nucleotides with the diluted phase, and simulate RNA polymerization inside FUS condensates with different cutoffs for bond formation. In Figure S6(a) we show the amount of free nucleotides left in the simulation box with different cutoff distances as indicated in the Figure legend. We find that a cutoff distance of 6.5 Å does not allow the formation of a single RNA-RNA bond due to residue-residue repulsion. Moreover, at cutoff distances 7.5 and 8.5 Å we find an almost identical rate of free nucleotides depletion. In light of this result, we opt to choose 7.5 Å due to the fact that, in our computational framework, the formation of a bond is performed dynamically during the course of the simulation, using the *fix*

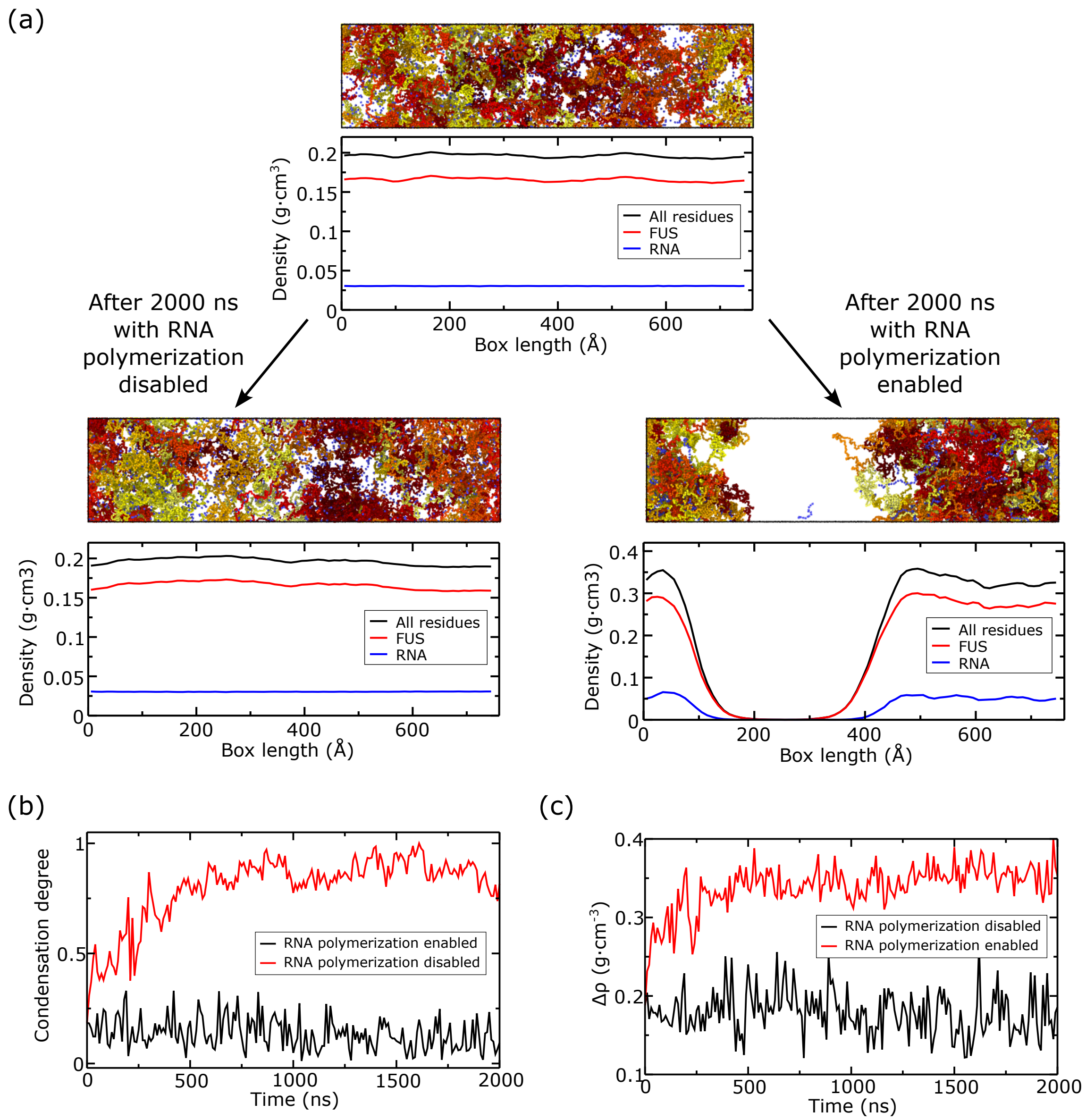


Figure S5: FUS/RNA simulation at the critical temperature (337 K) with constant number of molecules (no reservoir) and RNA polymerization enabled and disabled (control simulation) as indicated below. (a) Top: Rendered image of the initial configuration with all the FUS and RNA molecules dispersed, accompanied by the density profile (computed along the elongated box dimension) where the different colours depict the different components of the system. Bottom left: Rendered image and density profile after 2000 ns have been simulated with no RNA polymerization. Bottom right: Rendered image and density profile after 2000 ns have been simulated with RNA polymerization enabled. All the density profiles shown are averaged over 10 ns i.e. The initial one is computed between 0 and 10 ns of the trajectory, while those at the bottom are averaged between 1990 and 2000 ns of the trajectory. (b) Condensation degree as a function of time. (c) Density difference between the slabs with greater and lower density found in the simulation box (evaluated with the same slab criteria as the condensation degree).

*bond/react* command from LAMMPS as described in section SIV. Whenever a bond formation event takes place, effectively we insert a new harmonic bond with equilibrium distance 5 Å, as described by the Mpipi-recharged force field (section SI). As seen in Figure S6(a), the distance at which the two bond-forming residues will be between 6.5 (which is not reached in the test simulation we show in panel (a)) and 7.5 Å (the cutoff distance). Therefore, the bond formation event will necessarily generate a artificial potential energy increase, which will be dissipated over time. This potential energy discuntinuity is minimized when the residue-residue distance is closer to equilibrium, which happens when the cutoff distance is 7.5 Å. We

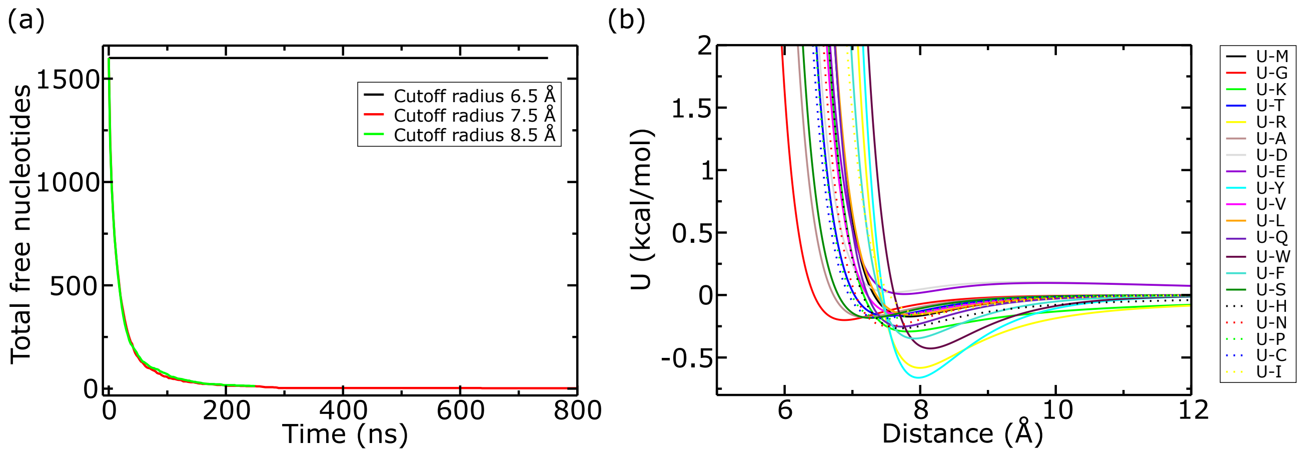


Figure S6: (a) Total amount of free uridine precursors present in a simulation with FUS at 287.5 K in absence of particle exchange between the diluted phase and a nucleotide reservoir, equivalent conditions to main text Figure 1. (b) Total potential energy of interaction between uracil and the different amino acids as a function of the inter-particle distance.

also conclude that this lower value is suitable given that even greater cutoff values are not required since, as seen in Figures 1-5 of the main text, this cutoff value allows us to observe and characterize the relevant condensation/dissolution events in a computationally affordable time. Therefore, 7.5 Å compromises computational accessibility with the least unphysical energy drifts and negligible dynamic observables difference with respect to greater and lower cutoff values.

Moreover, we set 10 Å as the distance cutoff at which RNA-protein contact is required in order to consider such nucleotide for a RNA-RNA bond formation. In Figure S6(b) we show the U-X (where X is the amino acid symbol) interaction potential for all uracil-amino acid pairs. At 10 Å we observe that the U-X contribution to the potential energy becomes minimal for all pairs, therefore defining a protein-crowded environment based on this distance criterion.

### SXIV. MED1 CONDENSATION/DISSOLUTION FURTHER CHARACTERIZATION

In this section we provide further characterization not included in the main Figures: In Figure S7(a) we characterize the time-dependent RNA content in the simulation. Moreover, Figure S7(b) represents the average RNA chain length as a function time. Lastly, Figure S7(c) shows different density profiles at different times that represent characteristic stages of the re-entrant MED1/RNA condensation, all of which averaged over 10 ns of simulation: First at the initial time, where RNA exists in low concentrations and short chain lengths, therefore not inducing condensation. Secondly, at $t = 400$ ns, where the MED1 condensate is formed once RNA has formed long enough chains to form stabilizing intermolecular contacts (see Figure 3c of the main text). Lastly, at $t = 800$ ns, when RNA is present in excess, condensation is no longer promoted, and the density profiles become flat once again, characteristic of a single dispersed phase. These 3 Figure panels characterize that the MED1/RNA re-entrant condensation process is accompanied by a gradual increase in the RNA content and chain length. Initial RNA growth favours condensation while posterior excess concentration leads to condensate dissolution. From Figure S7(a) it can also be concluded that it is at $t \simeq 300$ ns that a change in the rate at which the total amount of RNA residues increases takes place. This coincides with the consolidation of the MED1/RNA condensate (Figure 3(b) in the main text), and therefore the greater capacity to exchange RNA precursor particles with the reservoir. This addition of free nucleotides is also reflected in Figure S7(b) with a slight decrease in the average length, due to the addition of 1-len RNAs. Upon this RNA addition, a faster RNA elongation takes place (Figure S7(b) at $t > 500$ ns), rapidly leading to the condensate dissolution, linked to RNA over-accumulation. Once the condensate is disolved ($t > 600$ ns) we also observe a slight decrease in the total amount of RNA residues due to the excess present in the now single dispersed phase, which contains an RNA concentration greater than that of the reservoir.

[1] A. R. Tejedor, A. Aguirre Gonzalez, M. J. Maristany, P. Y. Chew, K. Russell, J. Ramirez, J. R. Espinosa, and R. Collepardo-Guevara, "Chemically informed coarse-graining of electrostatic forces in charge-rich biomolecular condensates," *ACS Central Science*, vol. 11, no. 2, pp. 302–321, 2025.

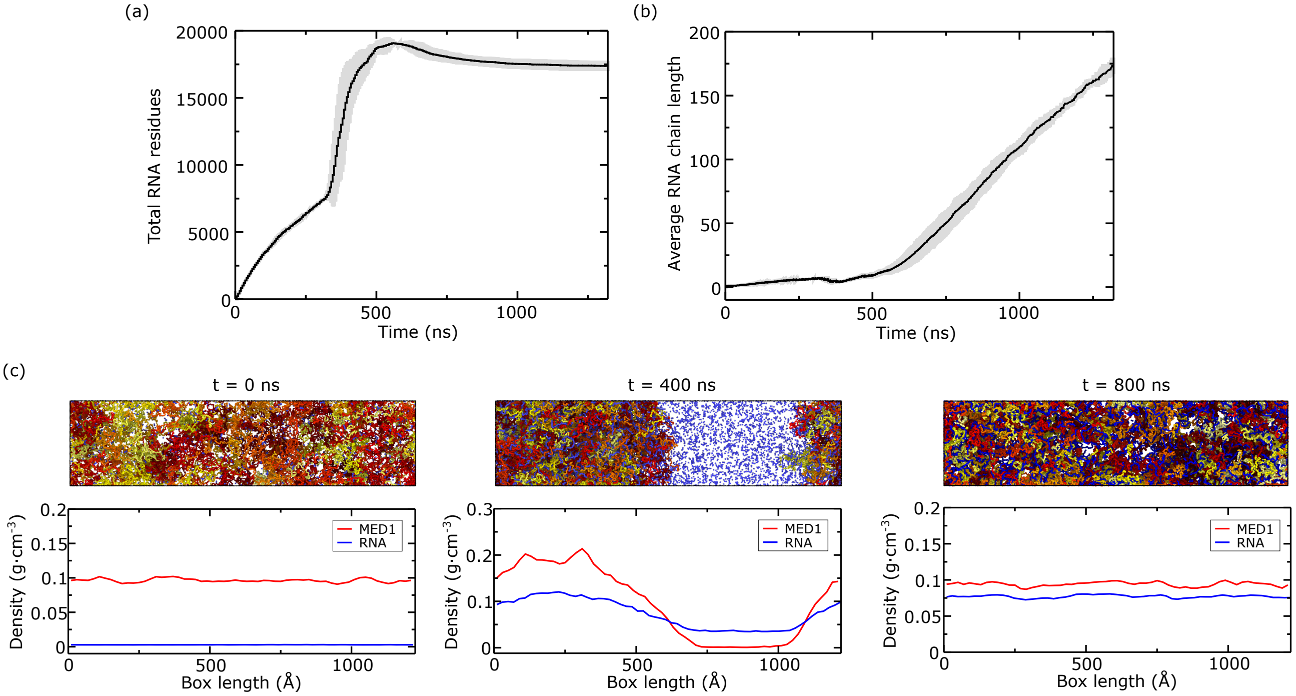


Figure S7: (a) RNA content as a function of time. (b) Average RNA chain length as a function time. (c) Density profiles for MED1 and RNA computed at 0 (left), 400 (middle) and 800 (right) ns. The density profiles were computed averaging 10 ns of simulation time, starting at the indicated time. On top of each density profile, a rendered image at the corresponding timestep is shown.

[12] J. S. Rowlinson and B. Widom, *Molecular theory of capillarity*. Courier Corporation, 2013.

[13] J. A. Zollweg and G. W. Mulholland, "On the law of the rectilinear diameter," *The Journal of Chemical Physics*, vol. 57, no. 3, pp. 1021–1025, 1972.

[14] A. Garaizar, J. R. Espinosa, J. A. Joseph, G. Krainer, Y. Shen, T. P. Knowles, and R. Collepardo-Guevara, "Aging can transform single-component protein condensates into multiphase architectures," *Proceedings of the National Academy of Sciences*, vol. 119, no. 26, p. e2119800119, 2022.

[15] A. Garaizar, J. R. Espinosa, J. A. Joseph, and R. Collepardo-Guevara, "Kinetic interplay between droplet maturation and coalescence modulates shape of aged protein condensates," *Scientific reports*, vol. 12, no. 1, p. 4390, 2022.

[16] A. R. Tejedor, I. Sanchez-Burgos, M. Estevez-Espinosa, A. Garaizar, R. Collepardo-Guevara, J. Ramirez, and J. R. Espinosa, "Protein structural transitions critically transform the network connectivity and viscoelasticity of rna-binding protein condensates

but rna can prevent it," *Nature communications*, vol. 13, no. 1, p. 5717, 2022.
